# A behavioral architecture for realistic simulations of *Drosophila* larva locomotion and foraging

**DOI:** 10.1101/2021.07.07.451470

**Authors:** Panagiotis Sakagiannis, Anna-Maria Jürgensen, Martin Paul Nawrot

**Affiliations:** Computational Systems Neuroscience, University of Cologne

## Abstract

The *Drosophila* larva is extensively used as model organism in neuroethological studies where precise behavioral tracking enables the statistical analysis of individual and population-level behavioral metrics that can inform mathematical models of larval behavior. Here, we propose a hierarchical model architecture comprising three layers to facilitate modular model construction, closed-loop simulations, and direct comparisons between empirical and simulated data. At the motor layer, the autonomous locomotory model is capable of performing exploration. Based on novel kinematic analyses our model features intermittent forward crawling that is phasically coupled to lateral bending. At the second layer, navigation is achieved via active sensing in a simulated environment and top-down modulation of locomotion. At the top layer, behavioral adaptation entails associative learning. We evaluate virtual larval behavior across agent-based simulations of autonomous free exploration, chemotaxis, and odor preference testing. Our behavioral architecture is ideally suited for the modular combination of neuromechanical, neural or mere statistical model components, facilitating their evaluation, comparison, extension and integration into multifunctional control architectures.

## Introduction

*Drosophila* larvae express a fairly tractable behavioral repertoire that is consistent across the 4-5 days of the larval life stage (***Almeida-Carvalho et al., 2017***). Most of the larval lifetime is dedicated to foraging for suitable nutrients while avoiding threats via stereotyped evasive behaviors. Behavioral control is achieved via neural circuits conserved throughout development (***Gerhard et al., 2017***), that span the entire tripartite CNS consisting of the brain, the subesophageal zone (SEZ) and the ventral nerve cord (VNC), making the larva a formidable system for studying control, execution and adaptation of behavior (***Zarin et al., 2019; Jovanic, 2020; Eschbach and Zlatic, 2020; Imambocus et al., 2022***). After reaching critical mass for pupation, homeostatic signals switch behavior towards food aversion, hypermobility and collaborative burrowing (***Wu et al., 2003***), terminating the feeding state and leading to pupation and metamorphosis.

High resolution methods for behavioral tracking (***Ohyama et al., 2013; Risse et al., 2017; Schumann and Triphan, 2020; Tadres and Louis, 2020; Thane et al., 2023; Laurent et al., 2024***), now routinely used in neuroethological experiments, have revealed detailed insight in the organisation of larval foraging behavior. In the absence of food resources, larvae engage in free exploration to locate food patches (***Sims et al., 2019***) with a characteristic alternation of locomotory activity and brief pauses (***Sakagiannis et al., 2020***), a property also reported for adult fly behavior (***Ueno et al., 2012; Reynolds et al., 2015***). Food consumption involves repetitive feeding motion and digging into the substrate (***Kim et al., 2017***). Statistical regularities that govern foraging behavior have been unveiled by analysis at the microscale of body kinematics and at the macroscale of larva trajectories (***Denisov et al., 2013; Risse et al., 2017; Karagyozov et al., 2018***). Locomotion combines the basic sensorimotor primitives of crawling and turning, and has been in the main focus of recent studies (***Heckscher et al., 2012; Wystrach et al., 2016; Sims et al., 2019***), whereas studies of feeding behavior remain scarce (***Ruiz-Dubreuil et al., 1996***). Both, crawling and feeding consist of recurring sensorimotor cycles controlled by central pattern generating circuits (***Heckscher et al., 2012; Mantziaris et al., 2020; Miroschnikow et al., 2018a***).

Salient olfactory cues can trigger appetitive or aversive chemotaxis, during which larvae navigate up or down a chemical gradient (***Gomez-Marin et al., 2011; Slater et al., 2015; Schleyer et al., 2015a; Klein et al., 2017***). During appetitive chemotaxis, the detection of minor concentration changes during lateral bending causes a directional bias in the turning movement, establishing a mechanism of active sensing (***Gomez-Marin and Louis, 2012; Wystrach et al., 2016; Thane et al., 2019***). Encounters with novel odorants in the presence of a food reward or a negative reinforcement such as the bitter taste substance quinine can induce olfactory learning, enabling short- and long-term behavioral adaptations (***Schleyer et al., 2011, Schleyer et al., 2015b; Gerber and Stocker, 2007; Diegelmann et al., 2013; Widmann et al., 2018; Weiglein et al., 2019; Jürgensen et al., 2024***). For quantification of choice behavior, widely-used standard group assays for behavioral preference testing have been established (***Gerber and Stocker, 2007; Gerber et al., 2014; Schleyer et al., 2015a,Schleyer et al., 2015b, Schleyer et al., 2011***).

The routine availability of detailed behavioral data and the broad experimental repertoire in neuroethology make the *Drosophila* larva an ideal model system for computational studies of behavioral control. Existing generative models typically address isolated aspects of larval behavior and vary widely in type and abstraction level, ranging from basic neuromechanics to abstract mathematical formulations.

To unify such diverse approaches, we introduce the concept of a *behavioral architecture*: a three-layered, hierarchical, and modular framework that organizes models from low-level behavioral primitives to high-level functions (***Figure 1A***). At the motor layer, we implement a refined coupled-oscillator model of larval locomotion capable of autonomous exploratory behavior, supported by a detailed kinematic analysis of larval tracking data during free exploration. Sensory modulation introduced at the intermediate (reactive) layer enables the model to reproduce key experimental findings of appetitive chemotaxis.

**Figure 1.**
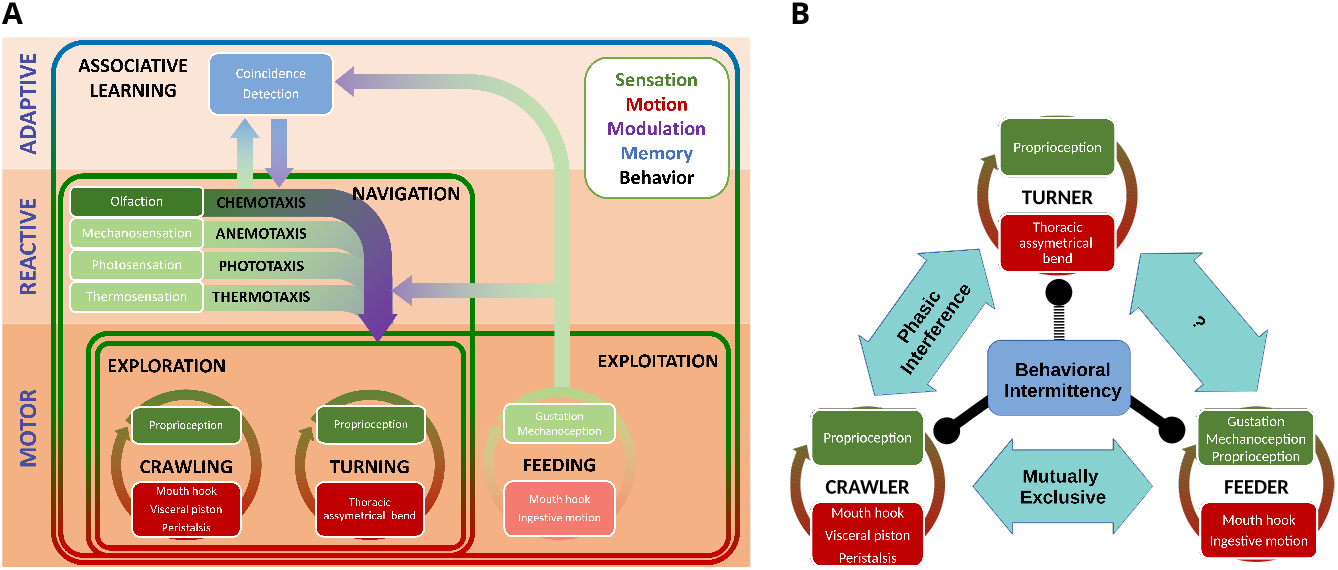
Behavioral architecture for larva foraging. **A:** In the three-layer hierarchical architecture the bottom layer (motor) consists of three basic sensorimotor effectors that constitute the locomotory model. The intermediate layer (reactive) features innate reactive behavior in response to sensory input that reflects changes in the environment. The top layer (adaptive) allows for behavioral adaptation through experience. Framed areas denote more complex behaviors that require subsumption of subordinate behaviors. The signal from feeding to the above layers is neuromodulatory, as indicated by the purple arrowhead.**B:** The intermittent coupled-oscillator locomotory model implementated at the motor layer. The turner and crawler module are phasically coupled during forward locomotion (head-casts and weathervaning are not separate modules; both emerge from this crawl–bend interference), while crawling and feeding are implemented as mutually exclusive sensorimotor primitives. Initiation or cessation of crawling is controlled by the intermittency module. Darker shades mark modules used in the present simulations; faded modules indicate possible extensions.

Finally, by combining the locomotory and sensory layers with a biologically realistic spiking neural network model of the mushroom body at the adaptive layer, the architecture reproduces empirical population-level learning scores across multi-trial associative conditioning protocols. Together, these modules allow agent-based, potentially closed-loop simulations of virtual larvae in virtual environments and generate behavior-level data comparable to experiments, facilitating model calibration, validation, and extension.

## Model

### Behavioral architecture

Larval behavior is hierarchically structured: basic sensorimotor primitives such as crawling, bending, and feeding can be combined into more complex functions such as exploration, taxis, or exploitation. A longstanding idea in neuroethology is that this behavioral hierarchy reflects a corresponding hierarchy of nested sensorimotor loops in the nervous system (***Prescott et al., 1999; Wilson and Prescott, 2022; Prescott and Wilson, 2023***). Within this view, low-level reflexive or rhythmic behaviors are generated locally by peripheral motor circuits, while more central brain areas recruit, modulate, or coordinate these primitives when more global or context-dependent integration is required. The subsumption architecture (***Brooks, 1986***) provides a functional modeling paradigm for such hierarchies: lower layers generate autonomous, stereotyped activity, while multisynaptic loops involving more centralized neural circuits act as descending modulators on the local circuits in order to coordinate global and complex behavior. The central idea is that energy-efficient decentralized neural control is the rule, while higher centers are recruited only when more extensive integration is needed e.g. in order to start, stop, accelerate, decelerate, or invert the activity of local loops (***Sen et al., 2017; Feng et al., 2020; Bidaye et al., 2020***). Importantly, the dimensionality of these descending pathways is much lower than the internal complexity of each layer. Layered control architectures have been widely used in behavior-based robotics (***Bicho, 1999; Prescott et al., 1999***) and offer a natural conceptual scaffold for integrating mechanistic, kinematic and abstract models of behavior. Here we adopt this paradigm as a simplifying organizing principle aligned with, rather than claiming to perfectly represent, the complete larval neuroanatomy.

We propose a three-layered behavioral architecture for *Drosophila* larval foraging (***Figure 1A***). The bottom, motor layer comprises the fundamental behaviors of crawling, turning, and feeding (***Figure 1B***). Each is implemented as a local sensorimotor loop between motor effectors and sensory feedback. Combinations of these primitives yield composite behaviors: for example, free exploration arises from integrating crawling and turning, and variation in their microstructure produces a continuum from local search to long-range dispersal. Consistent with this view, the ventral nerve cord (VNC) alone can sustain exploratory locomotion without descending input (***Sims et al., 2019***).

The intermediate layer integrates salient sensory cues across modalities and provides a unified descending modulation to the motor layer. This follows the subsumption principle of premotor convergence, where multisensory information is integrated into a common modulatory drive that shapes locomotion (***Wystrach et al., 2016; Eschbach and Zlatic, 2020***). Modulation of the motor layer by this single pathway enables reactive navigation along sensory gradients. In this study we focus on odor-driven chemotaxis, interpreted as an active sensing process in which larvae sample concentration changes during lateral bending to discover and ascend odor gradients. The same architectural principle generalizes to other modalities; for example, we have previously applied it to simulate active sensory sampling in thermotaxis across different *Drosophila* species (***Kafle et al., 2025***).

The top, adaptive layer implements associative learning, allowing larvae to modify innate stimulus valence and bias approach or avoidance decisions based on experience. The mushroom body is well established as the substrate for such learned valence in both larval and adult *Drosophila* (***Gerber and Stocker, 2007; Gerber et al., 2014; Schleyer et al., 2015a,Schleyer et al., 2015b, Schleyer et al., 2011; Owald and Waddell, 2015***). In line with these findings, the descending signal from the adaptive layer in our architecture is intentionally restricted to modulating approach versus avoidance tendencies, without altering lower-level motor primitives directly.

The behavioral architecture itself is agnostic to the level of abstraction or mechanistic detail within each module. Its purpose is to provide a modular framework in which diverse models - mechanistic, mathematical, or hybrid - can be positioned, interchanged, compared, or incrementally refined. To ensure compatibility despite differing internal complexities or time scales, every module exposes a clearly defined set of inputs and outputs. This design supports agent-based simulations of virtual larvae in virtual environments and allows seamless integration of mechanistic components as new empirical findings become available. To facilitate modeling within this framework, we provide the *larvaworld* Python package (https://pypi.org/project/larvaworld/), which implements the architecture, offers multiple instantiations of its modules, and enables direct comparison between simulated and experimental behavior.

### Locomotory model

We model larval locomotion in the two-dimensional plane by specifying forward velocity *v* and angular velocity ω as generated by crawling and bending, respectively. Building on previous oscillator-based approaches, our kinematic analysis motivates two key extensions: (i) intermittent crawling composed of runs and pauses, and (ii) stride-phase–dependent attenuation of angular motion. Together these yield an intermittent coupled-oscillator locomotory model (***Figure 1B***); its mathematical formulation is provided in the Materials and Methods, and the calibration pipeline in ***Appendix 2***.

To balance biological interpretability and computational tractability, we represent the body as two connected segments. The body state is therefore fully described by three variables: the midpoint position (***Appendix 1–Figure 1***), the front-segment orientation *θ*, and the bending angle *θ*_*b*_ between segments. This abstraction follows common practice in larval kinematics (***Gomez-Marin et al., 2011; Lahiri et al., 2011; Paisios et al., 2017***) and avoids the complexity of a full neurome-chanical model (see ‘Limitations of the study’). Crawling advances the midpoint along *θ*, whereas bending rotates the front segment; forward motion gradually restores *θ*_*b*_ toward zero by realigning the rear segment.

The locomotory model consists of two interacting oscillators. The crawler oscillator generates stride cycles that produce forward velocity *v*, with a tonic input *I*_*C*_ setting the crawling frequency *f*_*C*_ . The turner oscillator, based on ***Wystrach et al. (2016)***, generates alternating left–right bending with amplitude *A*_*T*_ . To capture the empirically observed crawl–bend interference, the bending amplitude is modulated by a Gaussian-shaped function of crawler phase, fitted to the data (red profile in ***Figure 3C***). In our implementation, weathervaning (continuous curvature changes during stridechains) and head casting (large reorientations during crawl-pauses) are not encoded as distinct modules. Instead, both arise from the same coupled-oscillator system, reducing model degrees of freedom while preserving the two canonical reorientation modes.

Intermittency is incorporated by alternating between crawling epochs (stridechains) and pauses, with stridechain length and pause duration sampled from empirical log-normal distributions (***Figure 3****B*). This stochastic state switching enables the model to reproduce the temporal organization of run–pause behavior observed in real larvae.

Within the subsumption-style architecture, higher layers can modulate the oscillators through only a few channels. For the crawler, top-down input may adjust the tonic drive *I*_*C*_ —thereby altering frequency—or initiate or terminate stride cycles. For the turner, top-down modulation of its tonic input affects both oscillation amplitude and frequency.

## Results

### Kinematic analysis of larval locomotion

To extract the core structure of larval locomotion, we analyzed experimental trajectories and body postures at both the single-animal (***Figure 2***) and population levels (***Figure 3***). These analyses reveal (i) the stride-based oscillation of forward velocity and the slower rhythm of lateral bending, (ii) a robust phase relationship between crawling and turning, and (iii) the intermittent organization of locomotion into stridechains and crawl-pauses.

**Figure 2.**
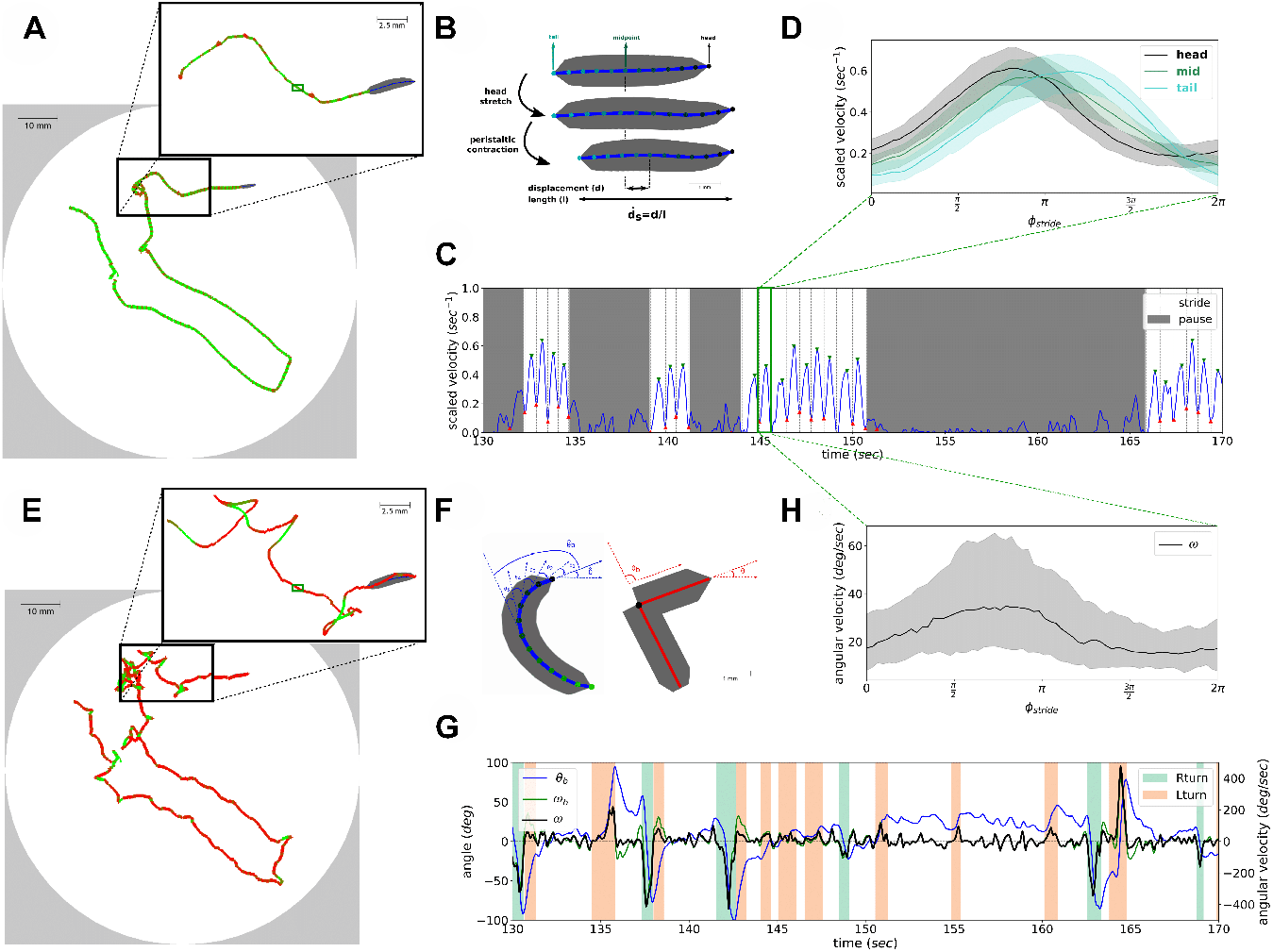
Kinematic analysis of a single Drosophila larva in locomotion. **A:** Individual larva trajectory tracking a posterior point along the midline of the animal . Trajectory color denotes the forward velocity *v* from 0 (red) to maximum (green). Inset focuses on a shorter track epoch analyzed in C and G. The full-length trajectory and the epoch in the inset are shown in ***Figure 2***—***video 1*** and ***Figure 2***—***video 2*** respectively. Dark green rectangle denotes the single stride described in B. **B:** Sketch of the single crawling stride indicated in A. The larva first stretches its head forward, anchors it to the substrate and then drags its body forward via peristaltic contraction. Scaled stride displacement 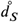 is defined as the resulting displacement *d* divided by the body-length *l*. **C:** Scaled forward velocity *v* during the 40 s track epoch selected in A (inset). Green and red markers denote the local maxima and minima used for stride annotation. Individual strides are tiled by vertical dashed lines. Successive strides constitute uninterrupted stridechains (white). Epochs that do not show any strides are annotated as crawl-pauses (gray). **D:** Scaled forward velocity *v* of head, midpoint and tail as a function of the stride cycle phase Φ. All detected strides have been interpolated to a stride oscillation cycle of period 2π. Solid lines denote the median, shaded areas the lower and upper quartiles across strides. **E:** Same trajectory as in (A) now tracking the head segment. Color denotes the absolute orientation angular velocity ω from 0 (red) to maximum (green). The full-length trajectory and the epoch in the inset are shown in ***Figure 2***—***video 1*** and ***Figure 2***—***video 2*** respectively. **F:** Definition of bending angle *θ*_*b*_ and orientation angle *θ* for the original 12-segment (blue) and the simplified 2-segment (red) larvae. **G:** Three angular parameters during the same track epoch shown in (C). Bending angle *θ*_*b*_, bend and orientation angular velocities ω_*b*_, ω are shown. Background shadings denote left and right turning bouts. For illustration purposes only turns resulting in a change of orientation angle Δ*θ* > 20^°^ are shown. **H:** Absolute orientation angular velocity ω during the stride cycle, as shown for *v* in (D). **Figure 2—video 1**. The full-length trajectory (A, E) colored according to angular and forward velocity. http://computational-systems-neuroscience.de/wp-content/uploads/2024/10/3.mp4 **Figure 2—video 2**. The short track epoch depicted in the insets (A, E) colored according to forward and angular velocity http://computational-systems-neuroscience.de/wp-content/uploads/2024/10/4.mp4

**Figure 3.**
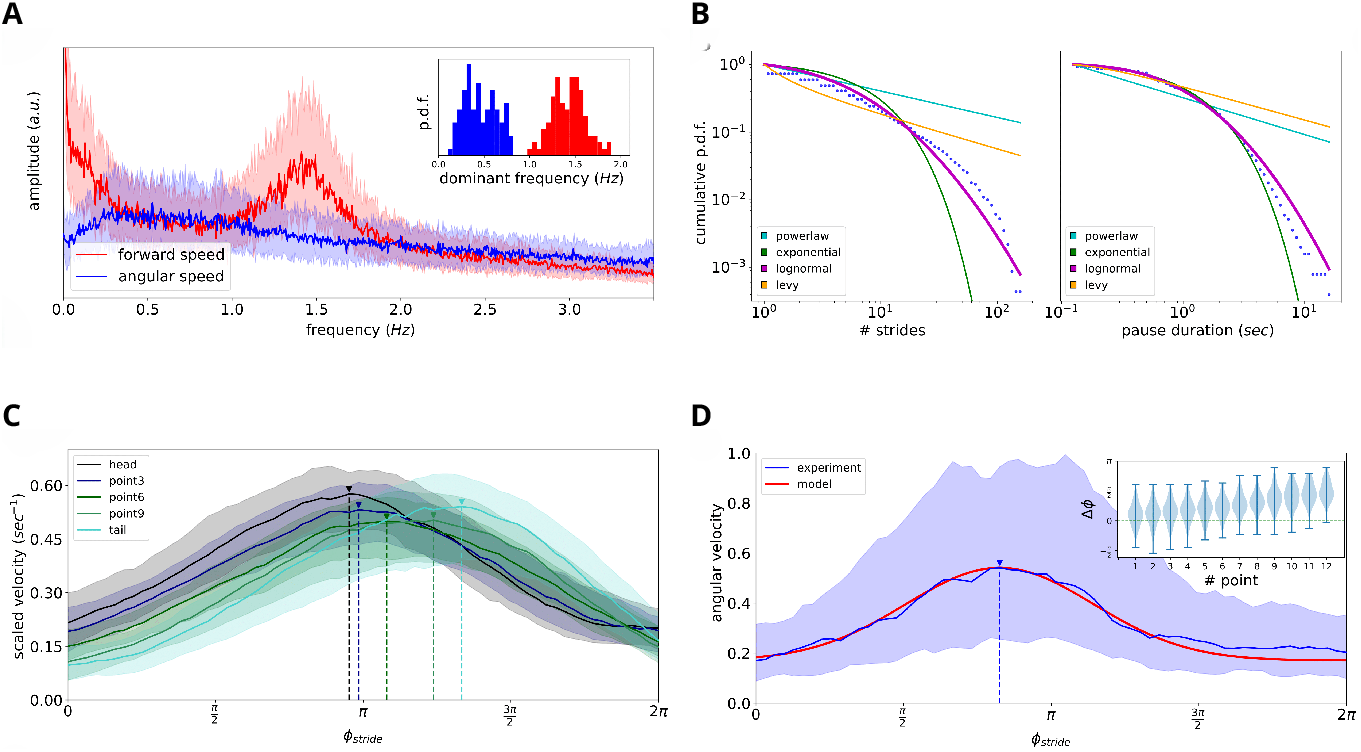
Population-level analysis. **A:** Fourier analysis of the forward *v* (red) and angular ω (blue) velocity across 100 larvae. Inset shows the respective dominant frequencies within suitable ranges 1 ⩽ *f*_*C*_ ⩽ 2.5 and 0.1 ⩽ *f*_*T*_ ⩽ 0.8 for *v* and ω, respectively. Crawling exhibits a dominant frequency of around 1.4 while lateral bending has a slower more variable rhythm of around 0.4. **B:** Epoch-duration distribution. Dots describe the cumulative probability density over logarithmic bins for the length of stridechains and the duration of crawl-pauses pooled across the larva population. Lines indicate the distribution with the lowest Kolmogorov-Smirnov distance among the best-fitting power-law, exponential, log-normal and Levy distributions. Stridechain length and pause duration are best approximated by log-normal distributions. **C-D:** Crawl-bend interference. The stride cycle kinematics are depicted for a single individual. All detected strides have been interpolated into a 64-bin oscillation cycle of period 2π. **C:** Forward velocity of 5 points along the larva midline. Velocity is scaled to the larva body-length. **D:** Absolute angular velocity ω (blue) normalized by the average value 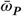 computed during the pause epochs. Fitted Gaussian function (red) describes well the phase-dependent attenuation imposed on ω and is used for the implementation of the coupled-oscillator locomotory model. Solid lines denote the median, shaded areas the lower and upper quartiles. Vertical dashed lines denote the cycle phase where the respective velocity reaches its maximum value. Inset : Phase offset Δ*ϕ* between the peak phase of each midline point’s forward velocity 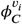 and the peak phase of angular velocity 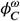 across a dataset of 100 tracked larvae. Notably, ω reaches its maximum just before the head forward velocity reaches its maximum.

### Single-larva kinematics

We first quantified translational and angular kinematics along individual trajectories. Tracking the forward velocity *v* of a posterior midline point (***Figure 2A***) shows that locomotion consists of consecutive peristaltic strides: *v* alternates between elevated (green) and reduced (red) values as the larva advances. A single stride is sketched in ***Figure 2B***, illustrating the extend–anchor–contract sequence whose amplitude is quantified by the scaled stride displacement 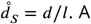 40-s epoch from the same trajectory (***Figure 2C***) shows stride maxima and minima (green and red) and how individual strides tile into stridechains (white) separated by crawl-pauses (gray). Videos of the full and inset trajectories are provided in ***Figure 2***—***video 1*** and ***Figure 2***—***video 2***.

To compare stride kinematics across occurrences, all detected strides were aligned to a normalized oscillation cycle of period 2π. The resulting forward-velocity profiles for head, midpoint, and tail segments (***Figure 2D***) reveal a stereotyped waveform consistent with a propagating peristaltic contraction.

Angular kinematics exhibit similar structure. Tracking the head’s angular velocity ω (***Figure 2E***) shows gradual reorientations during stridechains (weathervaning) and larger turns during crawl-pauses (headcasts). The bending geometry of the original 12-segment and simplified 2-segment representations is illustrated in ***Figure 2F***. During the same 40-s epoch, the bending angle *θ*_*b*_ and angular velocities ω_*b*_ and ω follow consistent temporal patterns (***Figure 2G***). Projecting ω onto the stride cycle (***Figure 2H***) reveals a smooth unimodal dependence peaking early in the cycle, indicating that turning amplitude varies systematically over the stride.

### Population-level rhythmicity, phase dependence, and intermittency

Across 100 larvae, Fourier analysis of forward and angular velocities (***Figure 3A***) reveal two dominant rhythms: a peristaltic crawling frequency around *f*_*C*_ ≈ 1.4 Hz and a slower bending rhythm around *f*_*T*_ ≈ 0.4 Hz, consistent with prior work (***Heckscher et al., 2012; Mantziaris et al., 2020; Thane et al., 2023; Wystrach et al., 2016***). These rhythms structure how larvae advance and reorient across time.

Larval crawling is intermittent, consisting of variable-length stridechains alternating with short crawl-pauses. Pooling stridechain lengths and pause durations across the dataset (***Figure 3B***) shows that both are best described by log-normal distributions among the candidate power-law, exponential, log-normal, and Lévy fits.

To quantify the phase relationship between crawling and bending, strides from all larvae were interpolated into a common 64-bin phase cycle. The progression of forward velocity from head to tail during the peristaltic cycle is illustrated in ***Figure 3C***. The angular velocity normalized by the mean pause-epoch value, 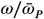, exhibits a clear stride-phase-dependent modulation (***Figure 3D***): bending is suppressed for most of the stride but briefly relieved early in the cycle. A Gaussian function (red) captures this unimodal attenuation profile and forms the basis for the phase-coupling mechanism incorporated in our locomotory model. The inset quantifies the phase offset Δ*ϕ* between the stride-phase peaks of forward and angular velocity across animals. Notably, the angular velocity peaks just before the head forward velocity reaches its maximum, suggesting a simple mechanistic explanation (see Discussion).

Together, the single-animal and population-level analyses reveal that larval locomotion is governed by a fast stride rhythm, a slower bending rhythm, a systematic phase relationship between them, and a stochastic alternation of runs and pauses. These empirical constraints directly motivate the intermittent coupled-oscillator model introduced below.

### Simulation of behavioral experiments

In this section we simulate increasingly complex behavioral experiments. We start with free exploration in the absence of navigation-guiding gradients, which does not imply absence of all sensory input but rather mirrors standard free-exploration lab assays. Then we advance to chemotactic navigation and finally to adaptive odor preference experiments. We simulate individual larvae using the model as calibrated in ***Appendix 2*** and all model parameters are shown in ***Appendix 2-Figure 1A***. Larval populations are constructed by pooling individual virtual larvae that behave independently as they move through the spatial arena and odorscape (***Niewalda et al., 2014***). In the present simulations, interlarval interactions are not modeled but rather each agent moves independently, consistent with the calibration datasets.

#### Exploration

In the first virtual experiment, we consider free exploration (***Figure 4***) and evaluate our simulation results against an empirical dataset (see Materials and Methods). To this end we simulate a population of 200 virtual larvae during three minutes while freely exploring a non-nutritious substrate in a Petri-dish, mimicking the experimental lab conditions (***Figure 4***). Because larvae typically reach the arena boundary within this time window, this experimental configuration only captures short-horizon exploratory dynamics. Consequently, the proposed intermittent coupled-oscillator locomotory model at the motor layer is sufficient to autonomously generate realistic exploration on these timescales, whereas assessment of long-horizon dispersal statistics reported in extended recordings (***Sims et al., 2019***) will require future simulations in larger or unbounded arenas.

**Figure 4.**
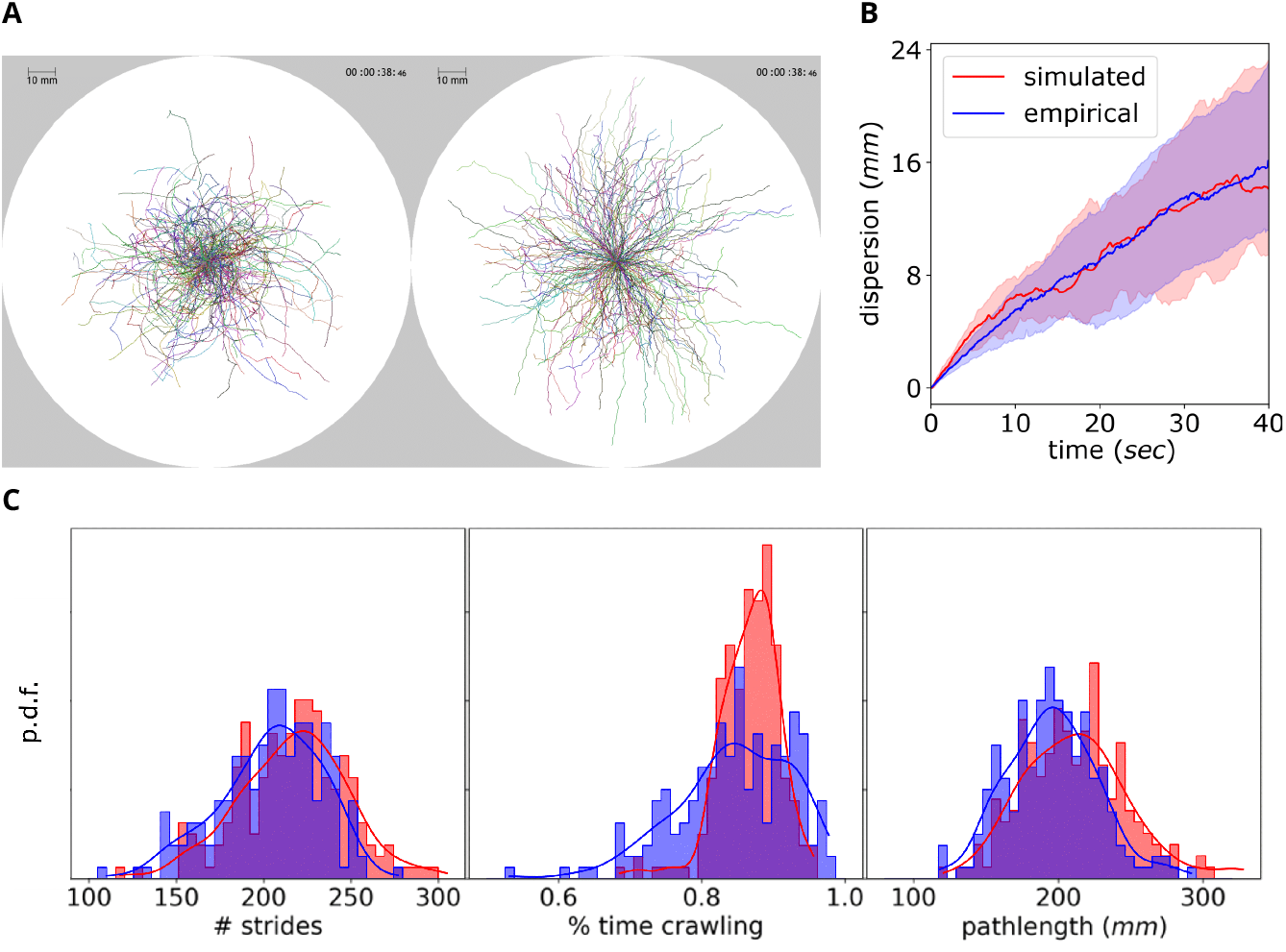
Free exploration in simulation and experiment. **A:** Dispersal of 200 larvae in experiment (left) and simulation (right) during 40 seconds. Individual tracks have been transposed to originate from the center of the arena. The entire temporal course shown in ***Figure 4***—***video 1*. B:** Dispersal from origin (Euclidean distance during the initial 40 s; larvae approach arena edges thereafter). Line indicates the group median while shaded area denotes first and third quartiles. **C:** Histograms for total number of strides, time ratio allocated to crawling and pathlength (cumulative distance). (arena dimensions = 500×500mm, N = 200 larvae, experiment duration = 3 minutes, simulation timestep =1/16). **Figure 4—video 1**. The temporal course of dispersal for real (left) and virtual (right) larvae shown in (A). http://computational-systems-neuroscience.de/wp-content/uploads/2024/10/5.mp4

Larvae are initially placed at the center of the dish, each with a random body orientation. Over the course of the experiment, both the simulated and the real larval population freely explores a petri-dish (***Video 1***) and disperses in space (***Figure 4A***, ***Figure 4***—***video 1***).

Among others, we report two complementary spatial metrics: radial dispersal (Euclidean distance from the origin at the end of the trajectory) and pathlength (cumulative distance traveled). Dispersal is shown during the initial 40 s because larvae typically reach the arena edge afterwards. The quantitative comparison shows good agreement between the simulated and empirical data with respect to the radial dispersal from the initial position (***Figure 4B***). Dispersal serves as an individual-level displacement metric from which population-level measures such as the mean-squared displacement (MSD) can be directly derived. Importantly, variability between individual animals is well captured by the variability between individual model instances, as shown for number of strides, crawling time, and pathlength (***Figure 4C***).

**Video 1. Free exploration simulation**. A population of 25 real (left) or virtual (right) larvae is placed on a dish and left to freely explore. The body of the real larvae has been simplified into 2 segments as described in ***Video 4***.

http://computational-systems-neuroscience.de/wp-content/uploads/2024/10/9.mp4

#### Chemotaxis

Chemotaxis describes the process of exploiting an odor gradient in space to locate an attractive or avoid a repelling odor source. An olfactory sensor (olfactor) placed at the front end of the virtual body enables active sensing during body bending and allows detection of concentration changes that modulate turning behavior by acting on the lateral oscillator (turner). Specifically, instantaneous odor sampling during lateral bending modulates its frequency and amplitude, while the peristaltic oscillator (crawler) remains unaffected, as proposed in (***Wystrach et al., 2016***).

To assess the chemotactic efficiency of our coupled-oscillator model we reconstruct the arena and odor landscape (odorscape) of two behavioral experiments described in (***Gomez-Marin et al., 2011***). In the first, larvae are placed on the left side of the arena facing to the right. An appetitive odor source is placed on the right side. The virtual larvae navigate up the odor gradient approaching the source (***Figure 5A,C,E***), reproducing the experimental observation in Fig.1 C in (***Gomez-Marin et al., 2011***). In the second, both the odor source and the virtual larvae are placed at the center of the arena. The larvae perform localized exploration, generating trajectories across and around the odor source. (***Figure 5B,D,F***), again replicating the observation in Fig.1 D in (***Gomez-Marin et al., 2011***). Two sample simulations can be seen in ***Figure 5***—***video 1***. These chemotaxis simulations are intended as proof-of-concept demonstrations rather than quantitative fits; see ‘Limitations of the study’ for a discussion. Note that all parameters of the motor layer (crawling, lateral bending, crawl-bend interference, and crawl-pause intermittency) were fitted on exploration data only and then kept fixed for chemotaxis and odor preference simulations.

**Figure 5.**
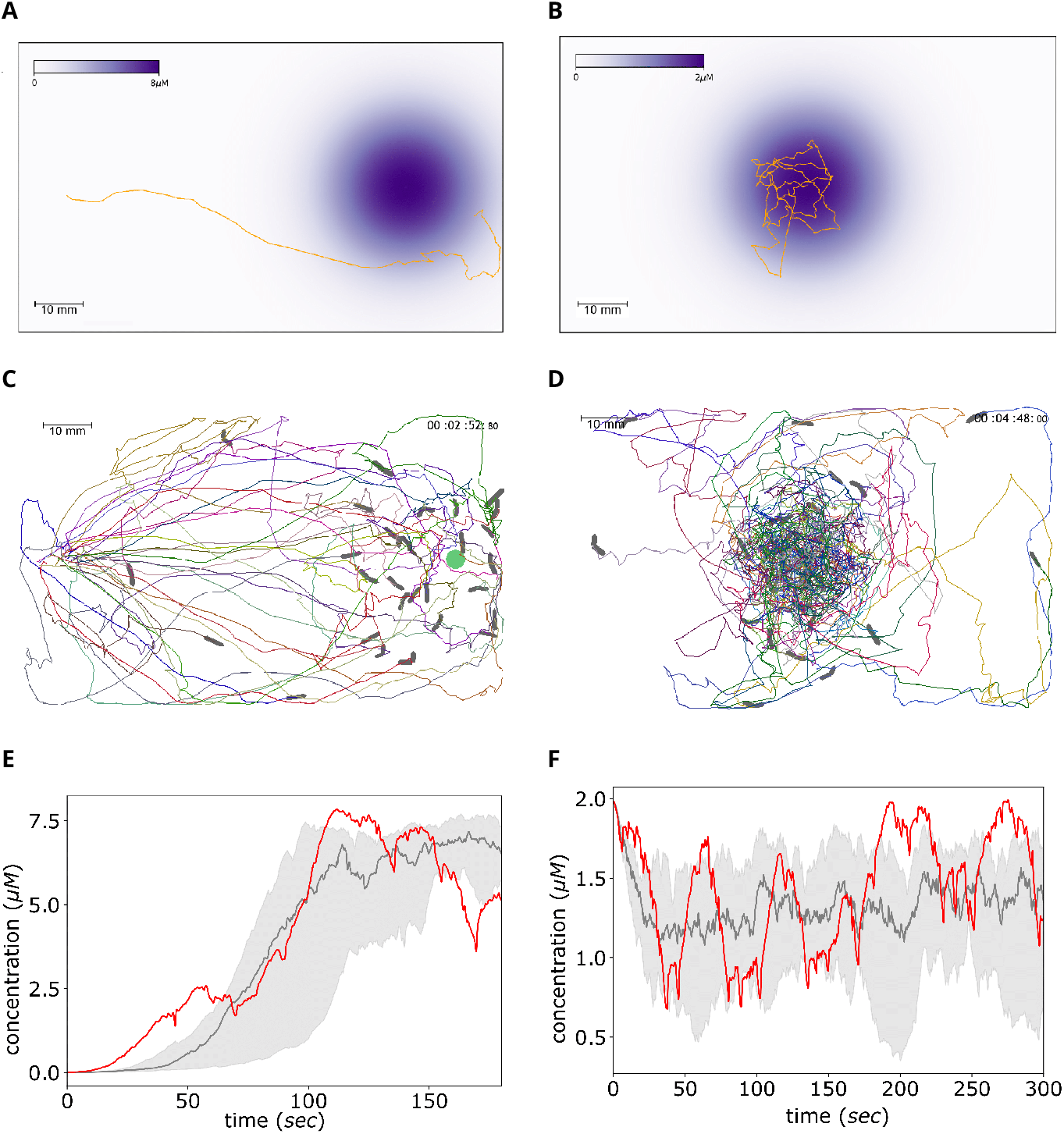
Simulation of chemotaxis. **A:** Experiment 1: A single odor source of 8.9*μM* peak concentration is placed on the right side of the rectangular arena creating a chemical gradient as indicated by the color scale. Larvae are placed on the left side facing to the right. Larvae are expected to navigate up the gradient approaching the source. A single larva trajectory is shown. This setup mimics the first experiment in ***Gomez-Marin et al. (2011)***. **B:** Experiment 2: A single odor source of 2.0*μM* peak concentration is placed at the center of the rectangular arena. Larvae are placed in close proximity to the odor source. Larvae are expected to locally explore generating trajectories around and across the source. A single larva trajectory is shown. This setup mimics the second experiment in ***Gomez-Marin et al. (2011)***. **C**,**D:** The trajectories of 25 virtual larvae during the two experiments. **E**,**F:** The odor concentration encountered by the virtual larvae as a function of time, measured in *μM*. Red curves refer to the single larva in A and B. Gray denotes the mean and quartiles of all 25 larvae in C and D. The simulation results fit well to the experimental estimates of concentration sensing during larval chemotaxis in ***Gomez-Marin et al. (2011)*** (arena dimensions = 100×60mm, N = 30 larvae, experiment duration = 3 and 5 minutes respectively, simulation timestep =1/16). Because the present chemotaxis implementation is a qualitative reproduction of the behavioral patterns reported by ***Gomez-Marin et al. (2011)*** and not a quantitatively calibrated model, we do not report success rates, approach distances, or track-length statistics; these metrics depend strongly on the specific odorscape geometry and experimental constraints and therefore lie outside the scope of the proof-of-concept simulations presented here.Parameterization of the motor layer was kept identical to exploration simulations. **Figure 5—video 1**. Time course of the two simulated chemotaxis experiments. http://computational-systems-neuroscience.de/wp-content/uploads/2024/10/6.mp4

#### Odor preference test

We simulated the standard odor preference assay described in the Maggot Learning Manual (***Michels et al., 2017***). Larvae were initialized at the center of a circular arena containing two odor sources placed on opposite sides. Each odor source was modeled as a Gaussian concentration field, producing partially overlapping attractive or aversive gradients. After three minutes of exploration, larval positions were used to compute the population-level Preference Index (PI) of the left odor,

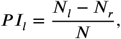

where *N*_*l*_ and *N*_*r*_ are the numbers of larvae on the left and right halves of the dish, respectively.

Because the turning modulation depends on an odor-specific gain *G*, expressed in arbitrary units, we first determined a suitable behavioral operating range. We performed a parameter sweep over 25^2^ combinations of left- and right-odor gains in *G* ∈ [−100, 100] using 30 simulated larvae per condition. The resulting PIs form the landscape shown in ***Figure 6A***. Example simulations for appetitive and aversive odors are shown in ***Figure 6***—***video 1***.

**Figure 6.**
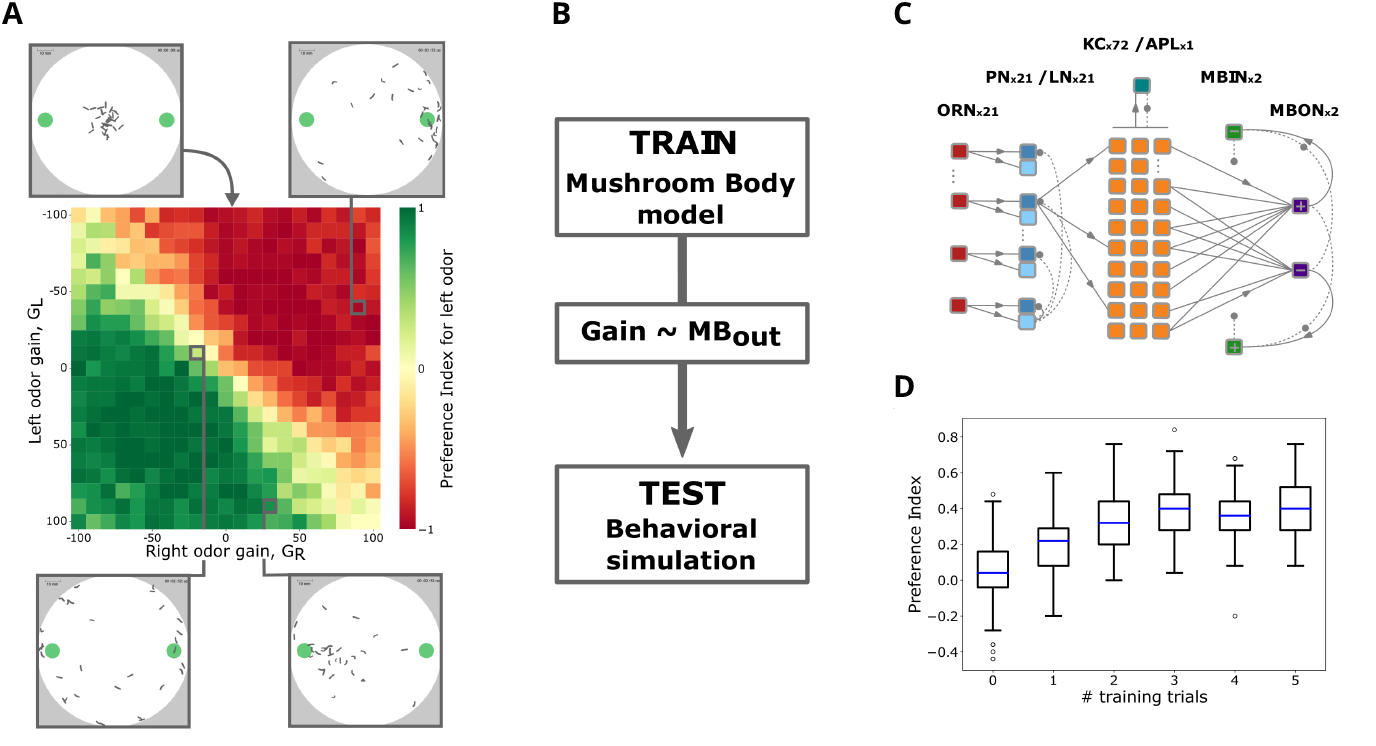
Simulation of innate and learned odor preference. **A:** A total of 25^2^ simulations are shown with the resulting Preference Index for different gains of the left and right odor. On the top left the initial state is shown with the larvae randomly generated at the center of the dish. The final state of three additional simulations is depicted on the top right and bottom left and right. See***Figure 6***—***video 1*** for videos of two sample simulations. **B:** The pipeline used for coupling the Mushroom Body (MB) model with the behavioral simulation. First a MB model is trained via a classical conditioning experiment where olfactory input is combined with reward. The resulting odor valence *MB*_*out*_ is then converted to odor gain *G* via a simple linear transformation and used to generate a virtual larva. Finally the odor preference of a virtual larva population is evaluated in a behavioral simulation. **C:** The spiking neural network comprising the MB model. The number of neurons comprising each layer is indicated. **D:** The resulting PIs for 100 simulations per number of training trials. In each of the 100 simulations per condition a population of 30 virtual larvae was generated and evaluated using a different random seed, always bearing the exact same 30 odor gains derived from the respective group of 30 trained MB models (arena dimensions = 100*x*100mm, N = 30 larvae, experiment duration = 3 minutes, simulation timestep =0.1. **Figure 6—video 1**. Two odor preference simulations, one with an appetitive and one with an aversive odor source placed on the left side of the dish. A non-valenced odor source is placed on the right side. http://computational-systems-neuroscience.de/wp-content/uploads/2024/10/7.mp4

To simulate learned odor preference, we coupled the behavioral simulations to the spiking mushroom body (MB) model of ***Jürgensen et al. (2024)*** (see Methods). In brief, the MB model was trained using the classical conditioning protocol of ***Michels et al. (2017)***, where an odor (CS^+^) was paired with sugar reward for five minutes. Five groups of 30 MB models underwent one to five conditioning trials (***Weiglein et al., 2019***). Each trained MB model returned an odor-valence signal *MB*_*out*_, which we converted to an odor gain *G* via a linear transformation (pipeline shown in ***Figure 6B***). This produced 30 odor gains per training condition, each used to generate one virtual larva.

Each virtual population of 30 larvae was then tested in an odor preference assay with the previously rewarded odor (CS^+^) on one side and a neutral odor on the other (***Figure 6***—***video 1***). To obtain robust population-level statistics, we repeated each condition 100 times with different random seeds, yielding 600 simulations in total. The resulting PIs (***Figure 6D***) increase with the number of conditioning trials and saturate after several pairings, in agreement with empirical observations (***Weiglein et al., 2019***).

Although group assays provide only population-level PIs and not individual trajectories, each simulated PI in ***Figure 6D*** arises from explicit larval movement dynamics. Thus, while no experimental single-larva trajectories are available for direct comparison, model-generated trajectories can be inspected on demand. The variability observed across the 100 repeats per condition originates solely from the behavioral simulation and resembles the variability seen across repeated real experiments.

The present implementation sequentially couples a trained MB model to the behavioral simulation. In the Discussion, we outline how a future closed-loop integration could enable full simulation of both conditioning and testing phases within a unified virtual environment.

## Discussion

### Experimental evidence for layered behavioral architecture

The behavioral architecture rests on two principles: (i) horizontal decomposition into functional modules that each generate a well-defined behavior, and (ii) vertical layering into semi-independent control levels, where higher layers modulat - rather than replace - the autonomous activity of lower ones. Here we summarize the neuroanatomical and behavioral findings that support this modular and hierarchical organization.

The basic larval behaviors captured at the motor layer (***Figure 1A***) have well-characterized neural substrates. Crawling consists of stereotyped repetitive strides, combining head–tail contractions driven by a “visceral piston” mechanism with a bilaterally symmetric peristaltic wave propagating from posterior to anterior segments (***Heckscher et al., 2012***) (***Figure 2B***). This motion is produced by segmental central pattern generators (CPGs) interconnected through short- and long-range premotor and interneuronal motifs (***Kohsaka et al., 2019; Zarin et al., 2019; Mantziaris et al., 2020***). Lateral bending arises from asymmetric activation of body-wall musculature initiated at thoracic segments (***Berni, 2015***). Feeding behavior is generated by mono- and multisynaptic sensorimotor loops linking enteric, pharyngeal, and external sensory organs to motor neurons controlling mouth-hook movement, head tilt, and pharyngeal pumping (***Miroschnikow et al., 2018b; Schoofs et al., 2024***). These circuits can operate without descending input: larvae continue to explore even after brain ablation (***Sims et al., 2019***). Because exploitation of food integrates locomotion with continuous ingestion and repositioning, we postulate that consummatory circuits in the subesophageal zone (SEZ) can sustain autonomous exploitation.

Higher brain centers modulate these basic behaviors through diverse neuromodulatory pathways. Dopamine, serotonin, acetylcholine, and octopamine play key roles at multiple levels of sensorimotor integration (***Berni et al., 2012; Zhang et al., 2013; Miroschnikow et al., 2018b; Malloy et al., 2019; Eschbach et al., 2020; Vogt et al., 2021***). Exploration–exploitation transitions can be acutely induced by dopaminergic signaling (***Schleyer et al., 2020***), while long-term balancing between the two states is mediated by hugin-dependent homeostatic neuromodulation (***Schoofs et al., 2014***). Anatomical mapping of sensory pathways to motor neuropils further highlights the contribution of interoceptive signals to behavioral regulation (***Qian et al., 2018***).

Olfactory modulation of locomotion also conforms to this layered view. Innate approach or avoidance to odors relies on a largely hard-wired pathway involving the antennal lobe (AL) and its projection to the lateral horn (LH), in both larvae (***Schulze et al., 2015; Vogt et al., 2021***) and adults (***de Belle and Heisenberg, 1994; Strutz et al., 2014; Dolan et al., 2018***). Learned odor valence, by contrast, strongly depends on descending control from the mushroom body (MB) in larvae (***Saumweber et al., 2018; Eschbach et al., 2020***) and adults (***Slater et al., 2015***). These innate and learned pathways converge onto premotor networks downstream of the AL (***Schleyer et al., 2015a; Eschbach et al., 2020***), where they shape locomotory decisions. For example, LH-dependent descending input can transiently suppress crawling to trigger larger reorientation events when navigating down-gradient, aiding chemotaxis (***Tastekin et al., 2018***). Finally, the internal homeostatic state (e.g. starvation vs. satiation) modulates behavior via transmitter release and neuropeptide signaling at multiple processing stages—including AL, LH, and MB—linking metabolic needs to sensorimotor control (***Vogt et al., 2021; de Tredern et al., 2025***).

### Previous computational models of larva locomotion

Larva locomotion has been modeled extensively. Generative models of peristaltic crawling feature neural and/or neuromuscular dynamics. Sequential neural activity patterns across the segmental body can be generated by longitudinally repeated CPGs of paired excitatory and inhibitory population rate units, possibly involving proprioceptive feedback (***Gjorgjieva et al., 2013; Pehlevan et al., 2016***). This relatively abstract CPG model has been elaborated into a connectome-based circuitry of premotor and motor neurons. Under tonic activation the model exhibits two functionally distinct, though structurally overlapping, recurring patterns of neuromuscular activity, responsible for forward and backward peristalsis respectively (***Zarin et al., 2019***). At the other end of the modeling spectrum the contribution of the visceral-piston mechanism to the peristaltic cycle has been assessed in a biomechanical model (***Ross et al., 2015***). A more elaborate neuromechanical model, based on segmental localized reflexes and substrate frictional forces and assuming empirically informed axial and transverse oscillatory frequencies, has been shown to generate forward and backward crawling without the need for any neural activation pattern (***Loveless et al., 2019***).

Lateral bending has also been captured in statistical (***Davies et al., 2015***) or generative models (***Wystrach et al., 2016; Loveless et al., 2019; Loveless and Webb, 2018***). In the context of free-exploration it has been shown to be a byproduct of chaotic body neuromechanics underlying peristalsis (***Loveless et al., 2019***). In the context of chemotaxis it has been attributed to a distinct oscillatory process, autonomous (***Wystrach et al., 2016***) or semi-autonomous to crawling (***Davies et al., 2015***). In the former case oscillation is driven by mutual inhibition between excitatory-inhibitory circuits. In the latter, bending behavior is dissected into low-amplitude weathervaning while crawling and high-amplitude headcasting during crawl-pauses, an approach essentially equivalent to an attenuation of the lateral oscillation amplitude due to crawling. Autonomous exploration and chemotaxis can be generated by an integration of the neuromechanical and the independent lateral oscillator models (***Loveless and Webb, 2018***).

Apart from the rare occasion where such models implement mutually exclusive mechanistic hypotheses and are indubitably incompatible to each other, in most cases they are indeed complementary, overlapping, nested or disconnected in terms of the generative mechanisms they aim to capture and could potentially coexist under a broader control system. In this context our unifying architecture for larval behavior in which any partial model can be positioned can be valuable for modelers and roboticists, interested in a behavior-based synthetic approach.

Our design deliberately trades segmental neuromechanical detail for computational efficiency and modularity in the motor layer. The framework is explicitly designed so that more detailed neuromechanical or connectome-derived modules can replace the current ones without altering inter-layer contracts.

### Intermittent coupled-oscillator model for realistic locomotion

Each of the above models could be adjusted so that at minimum it generates the 2D translational and angular motion of a virtual body and could therefore populate the motor layer of the behavioral architecture. In this study we propose such a model, assembled in a synthetic approach by distinct modules, which either extend previously proposed locomotory models or integrate novel findings derived from our analysis of kinematic parameters. Given a simplified bisegmental body, a simple oscillator under frequency-regulating tonic activation is adequate for generating recurrent strides, efficiently summarizing the complex underlying neural and neuromechanical dynamics (***Heckscher et al., 2012; Mantziaris et al., 2020***). Concerning angular motion, the previously introduced lateral oscillator model (***Wystrach et al., 2016***) meets the requirements and can therefore be coupled to the forward oscillator.

Our proposed model contributes two novel features. First, the intermittent nature of crawling as transitions between runs and pauses (***Figure 3B***). And, second, the peristaltic cycle-phase dependent attenuation of angular motion (***Figure 3D***). By combining these two features, the two bending behavioral modes termed weathervaning and head-casting are naturally generated via a phasic coupling between the two oscillators.

#### Behavioral intermittency

Larval locomotion is intermittent meaning that crawling runs are transiently intermitted by brief pauses. Transitions between these two behavioral states occur autonomously during free exploration. Traditionally, in the context of movement ecology, intermittency has been studied in the spatial regime by characterizing the distributions of run distances and turn amplitudes occurring during brief stationary reorientation events (pauses). Power-law distributed runs in line with Levy-walk theoretical models (***Günther et al., 2016; Sims et al., 2019***) and diffusion-like kinematics have been reported (***Klein et al., 2017***). On the other hand, the turn amplitude distribution diverges from the original Levy-walk-predicted uniform distribution, as it is highly skewed towards small amplitudes, even if only significant reorientation events are taken into account (***Sims et al., 2019***). Moreover, the speed-curvature power-law relationship has been disputed (***Zago et al., 2016; Marken and Shaffer, 2017; Zago et al., 2017; Marken and Shaffer, 2018***). Regarding the temporal dynamics of intermittency, the duration distribution of these pauses is commonly neglected in traditional Levy-walk literature as they are usually characterised via the amplitude distribution of their resulting turning events. A recent analysis of larval tracks reported that the duration of pauses follows a power-law while that of activity bouts a log-normal distribution (***Sakagiannis et al., 2020***), partly in line with findings in adult-fly studies (***Ueno et al., 2012; Reynolds et al., 2015***). In our current study we found log-normal best fits for both pause duration and stridechain length. The disagreement over the pause duration might be attributed to the different timescale assessed in the two studies. Contrary to the high-resolution 180-sec tracks in this study, those analyzed previously lasted at minimum 1024 sec and up to 1 hour, allowing for the detection of longer pauses and imposing the necessity to fit over 4 orders of magnitude bypassing the apparent drop around 10 sec, also seen in Fig.3 of ***Sakagiannis et al. (2020)***.

Concerning computational modeling, the contribution of behavioral intermittency to locomotion has not been adequately appreciated. Candidate generative models can either simply sample statistical distributions (***Sims et al., 2019***) or feature a generative mechanism that yields state transitions. Stochastic state-transitions have been included in a model of larva exploration yielding exponentially distributed epochs of both runs and stationary headcasts (***Davies et al., 2015***). At a mechanistic level, a recent study presented a simple binary-neuron model exhibiting state transitions between power-law and non power-law regimes via self-limiting neuronal avalanches and proposed a plausible underlying mechanism that explains initiation/cessation of crawling (***Sakagiannis et al., 2020***). All these attempts can be considered instantiations of a behavioral intermittency module (***Figure 1B***) controlling cessation and re-initiation of crawling, central for generating epoch transitions. In the present study, we chose stochastic sampling from the empirically fitted parametric distribution models.

#### Crawl-bend interference

Crawling includes mouth hook motion. Specifically, the first phase of a crawling stride consists of concurrent forward motion of head and tail segments, aided by a ‘visceral pistoning’ mechanism that generates forward displacement of the gut. Subsequently, the mouth hooks anchor the head to the substrate so that the second phase of peristaltic motion can drag all other segments forward as well, completing the stride (***Heckscher et al., 2012***). With respect to turning, it is still debated whether individual turns should be considered as discrete reorientation events that are temporally non-overlapping with crawling bouts (***Sims et al., 2019***), or whether lateral bending occurs in an oscillatory fashion generating turns both during crawling (weathervaning) and during pauses (headcasts) (***Wystrach et al., 2016; Thane et al., 2019***). The latter is supported by detailed eigen-shape analysis confirming that larvae rarely crawl straight, rather forward locomotion is always accompanied by continuous small amplitude lateral bending (***Szigeti et al., 2015***). It follows that crawling does not exclude bending, rather the two strictly co-occur.

Crawling and bending partially recruit the same effector neural circuitry and body musculature at least at the level of the thorax. Peristaltic motion during crawling includes sequential symmetric bilateral contraction of all segments while bending occurs due to asymmetric unilateral contraction of the thoracic segments. This partial effector overlap could result in interference between the two processes. Indeed here we report a phase-dependent attenuation of angular velocity (***Figure 3C***). Attenuation reaches a minimum at a specific phase of the cycle, closely preceding the increase of head forward velocity (***Figure 3C***:*inset*). This coincides with the stride phase when the head stops being anchored to the substrate and is therefore free to move laterally. When applying a phase-dependent Gaussian attenuation kernel on angular velocity we managed to accurately reproduce the empirical relation (***Appendix 2-Figure 1C***).

A reasonable hypothesis would then be that the asymmetric thoracic contraction generating lateral bending becomes easier when the head is not anchored to the substrate therefore during a specific phase interval of the stride cycle. We propose that crawling phasically interferes with lateral bending because of these bodily constraints. A consequence of this hypothesis is that the amplitude of turns generated during crawl-pauses (headcasts) is larger in comparison to those generated during crawling (weathervaning) because during pauses the crawling interference to lateral bending is lifted. It is this phenomenon that dominates the description of larva exploration as a Levy-walk with non-overlapping straight runs and reorientation events, where weathervaning is neglected (***Günther et al., 2016; Sims et al., 2019***). Nevertheless, it has been included in a previous stochastic model of larva exploration, where it has been treated as entirely distinct to headcasts, occurring during crawl-pauses, via the application of differential constraints on both the angular velocity and the resulting turn amplitude (***Davies et al., 2015***). By implementing a coupled-oscillator locomotory model we avoid such dual treatment of headcasts and weathervaning.

### Olfactory learning in closed loop behavioral simulations

We have integrated a previously developed biologically realistic spiking MB model to demonstrate modularity: models with very different abstraction levels can be combined under the same architecture. This is not a bias toward modeling depth of the MB per se, but a stress-test of architectural flexibility. We have reproduced the results of a basic associative learning experiment in the fruit fly larva (***Schleyer et al., 2018; Jürgensen et al., 2024***) by the open-loop simulation of classical conditioning trials and subsequent closed-loop behavioral simulation during the memory retention test for individual virtual larvae (***Figure 6****B-D*). This modeling approach can be extended in multiple ways. First, the larva demonstrates a number of additional learning capabilities such as differential conditioning (***Schleyer et al., 2011, Schleyer et al., 2015b, Schleyer et al., 2018***), extinction learning (***Felsenberg and Waddell, 2019; Lesar et al., 2021***), and relief learning (***Saumweber et al., 2018; Weiglein et al., 2019; Gerber et al., 2014; König et al., 2018***). Interfacing neural network simulations with candidate circuit and synaptic mechanisms of plasticity with our behavioral model allows to directly compare virtual and empirical behavioral experiments, both at the level of individual and group assays. Second, while information about odor concentration is provided through active sensing, simultaneous input from a feeder module could activate the dopaminergic reward pathway required for synaptic plasticity in the mushroom body. At the same time, simulation of food uptake and energy expenditure will regulate the agent’s energy homeostasis. This would further allow realistic foraging scenarios with food depletion and competition among animals.

Closing the loop from active sensing to associative memory formation and behavioral control requires to synchronize a (spiking) neural network at the adaptive layer with the sensory module (reactive layer), and the locomotory and feeding modules at the motor layer (***Figure 1****A*). This will enable the simulation of virtual larvae experiencing spatial and temporal dynamics in a virtual environment or on a robotic platform (***Helgadóttir et al., 2013***), and it will allow to test model hypotheses on sensory-motor integration and to infer predictions for experimental interventions such as optogenetic stimulation (***Saumweber et al., 2018***) or genetic manipulation (***Saumweber et al., 2011; Michels et al., 2011; Widmann et al., 2016; Springer and Nawrot, 2021***).

### Limitations of the study

Several limitations mentioned throughout the study are summarized here : (i) The crawl–bend interference is statistically implemented, based on the kinematic analysis, without a complete segmental neuromechanical derivation. (ii) The intermittency module uses sampling from data-fitted stridechain and pause distributions, without an explicit neural switch model. (iii) Chemotaxis simulations are proof-of-concept and qualitatively fitted, not quantitatively calibrated to any published datasets. (iv) The intermittent coupled-oscillator model presented here does not attempt to cover the full larval repertoire, such as hunching, digging, backward crawling, or rolling, nor does it explicitly implement handedness as a directional bias. (v) Larva–larva interactions are not modeled here, nonetheless the *larvaworld* software supports tactile sensors for future interaction studies.

Our framework emphasizes modularity and computational tractability, which necessarily imposes several limitations. First, the locomotory model operates at a coarse kinematic level. Crawl–bend interference is implemented as a fitted, phase-dependent attenuation rather than being derived from segmental neuromechanics, and the intermittency module samples stridechains and pauses from empirical distributions without modeling the underlying neural mechanisms. Furthermore, behaviors requiring detailed musculature or segment-specific coordination (e.g. backward crawling, rolling, digging) are not modeled and long-term exploration statistics are not considered, nor is interindividual variability in turning preference (handedness).

Second, the navigation layer provides qualitative demonstrations of chemotaxis but is not quantitatively calibrated to odor-guided trajectories. Odor-driven modulation acts through a single pathway to the turning oscillator, and weathervaning and headcasts emerge from the same mechanism, which may under-represent the diversity of reorientation strategies. Third, at the adaptive layer, the mushroom body model is coupled to behavior sequentially rather than in closed loop: learning is performed independently of movement, and only the resulting odor valence is passed to the behavioral simulation.

Finally, the present study focuses on solitary larvae. Larva–larva interactions are not modeled explicitly, although the *larvaworld* framework supports tactile sensors for future interaction studies. These simplifications reflect deliberate design choices that maintain accessibility while allowing more mechanistic modules to be integrated in subsequent work.

## Materials and Methods

We first describe the experimental dataset and the software package used in this study. Then we explicitly describe each of the computational modules that comprise the proposed behavioral architecture. The metrics used throughout the analyses of both real and simulated datasets are described in ***Appendix 1***.

### Dataset description

The larva-tracking dataset was acquired by M. Schleyer and J. Thoener at the Leipzig Institute of Neurobiology. It comprises 31 experimental groups, each containing approximately 30 third-instar *Drosophila* larvae (Canton S). Larvae were placed on agarose-filled Petri dishes of 15 cm diameter without any added stimuli and video-recorded from above at 16 Hz for 3 minutes.

During tracking, 12 points were detected along the longitudinal body axis of each larva. For the present study, all groups were pooled into a single population, and timepoints involving detected collisions were removed. From this cleaned dataset, we selected the 200 most complete, uninterrupted trajectories for subsequent analysis.

The recorded *x*–*y* coordinate time series were smoothed with a first-order Butterworth low-pass filter (cutoff frequency 2 Hz) to suppress tracking-related noise while preserving the behaviorally relevant crawling frequency of *f*_*C*_ ≃ 1.4 Hz. The effects of insufficient and excessive filtering are illustrated in ***Video 2***.

**Video 2. Filter selection**. The effect of inadequate or excessive filtering of the empirical larva recordings is illustrated. The left video shows the jittery original recording while the effect of lowpass filtering at cut-off frequencies of 4 Hz, 2 Hz and 0.5 Hz is shown on the rest. Selection of an intermediate 2 Hz cut-off frequency eliminates the unrealistic jitter while preserving the behaviorally relevant crawling frequency.

http://computational-systems-neuroscience.de/wp-content/uploads/2024/10/8.mp4

### Software package and code availability

All data processing, analysis, and model simulations were performed using our open-source Python package *Larvaworld* (https://pypi.org/project/larvaworld/), a unified platform for behavioral analysis and simulation of *Drosophila* larvae. In *Larvaworld*, empirical and simulated data are handled identically: the same analysis pipeline and behavioral metrics are applied to both, ensuring methodological consistency throughout.

The behavioral architecture introduced in this manuscript provides the backbone for constructing, extending, configuring, and fitting behavioral models within *Larvaworld*. The intermittent coupled-oscillator model developed here is included, alongside other preconfigured models.

A separate publication describes the software architecture and usage in detail, and provides practical tutorials for science and teaching (***Sakagiannis et al., 2025***). Comprehensive documentation is available at https://larvaworld.readthedocs.io.

### Inter-individual variability in virtual populations

Traditional approaches in computational ethology typically use generative models tuned to reproduce the behavior of an *average* animal, thereby neglecting the substantial inter-individual variability observed within real populations. Here, we adopt a complementary, population-level approach that explicitly preserves individuality. Instead of instantiating a single representative model larva, we simulate a group of non-identical animats whose parameters are sampled from a fitted multivariate normal distribution that captures the empirical variability and pairwise correlations between parameters.

To construct this distribution, we quantified several endpoint behavioral and morphological parameters for 200 larvae and fitted a multivariate Gaussian model to the resulting measurements. A subset of three representative parameters—body length *l*, crawling frequency *f*_*C*_, and mean scaled stride displacement ^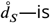^ —is shown in ***Figure 7***. Empirical univariate and bivariate distributions (blue) closely match the projections of the fitted multivariate Gaussian (red), which provides a compact generative model for individual variability. We further note that handedness represents an additional axis of individual variation in *Drosophila* larvae (***Wosniack et al., 2022***) and may be incorporated in future work.

**Figure 7.**
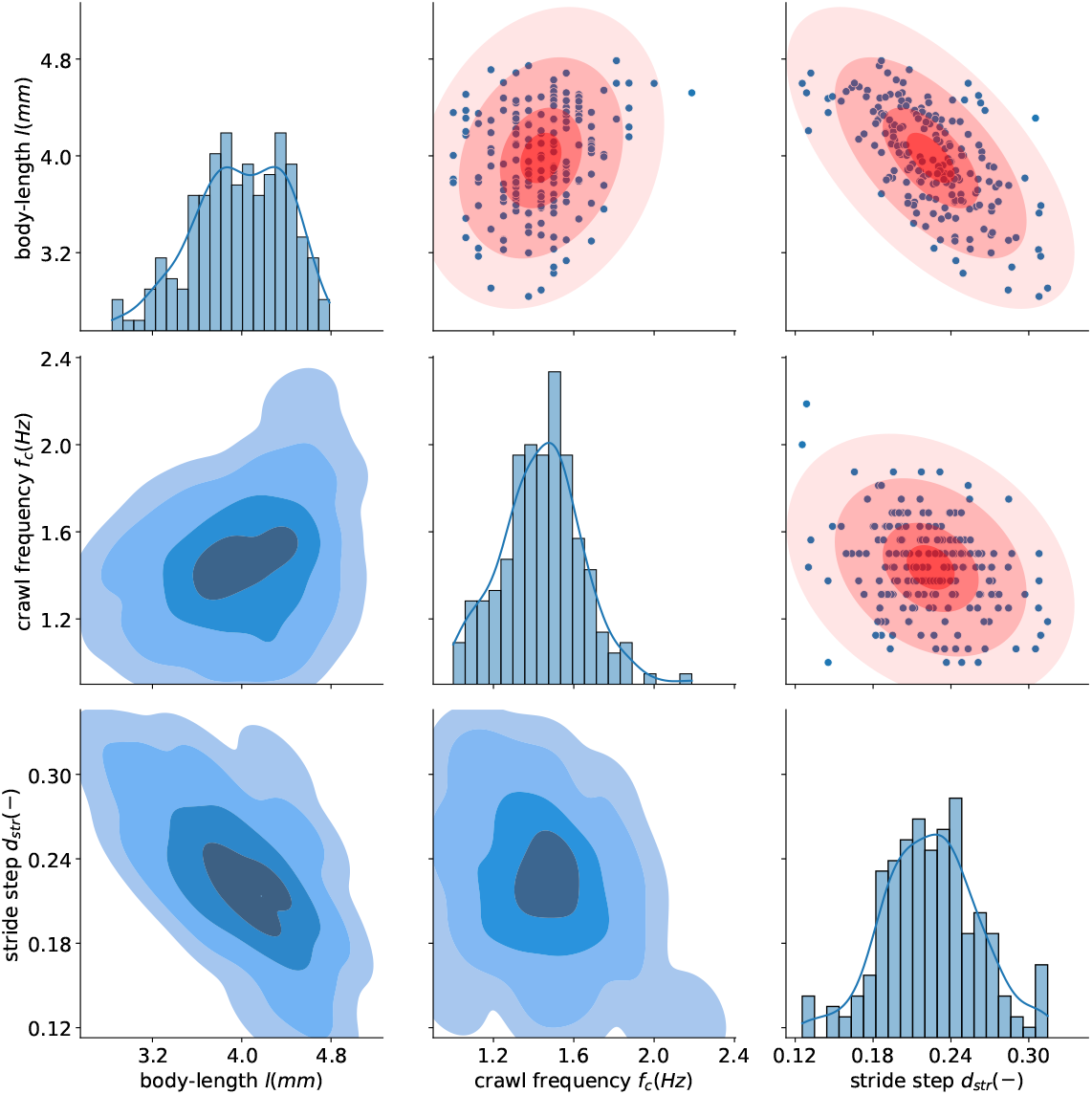
Individuality: empirical (blue) and fitted (red) parameter distributions. Diagonal panels show histogram and kernel density estimates (KDEs) for body length *l*, crawling frequency *f*_*C*_, and mean scaled displacement per stride 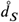 across a population of 200 larvae in the experimental dataset. Lower off-diagonal panels depict bivariate projections of the empirical three-dimensional KDE. Upper off-diagonal panels show the corresponding projections of the fitted multivariate Gaussian distribution, with ellipses indicating 0.5, 1, 2, and 3 standard deviations. In the virtual population, parameter sets for individual larvae are generated by sampling from this multivariate Gaussian. Blue dots represent empirically measured larvae.

### Module definition

The building blocks comprising the behavioral architecture are described as separate modules of defined input and output. The modular architecture only specifies the required placeholders and remains agnostic to the specific module implementations. Nevertheless, in what follows, the specific modular implementation for the proposed intermittent coupled-oscillator locomotory model will be described alongside the general module-placeholder definition. The parameters of the final model configuration are shown in ***Table 1*** and its modular function is illustrated in ***Video 3***. Calibration of each module on the empirical dataset and parameter specification is described in detail in ***Appendix 2***.

**Table 1.**
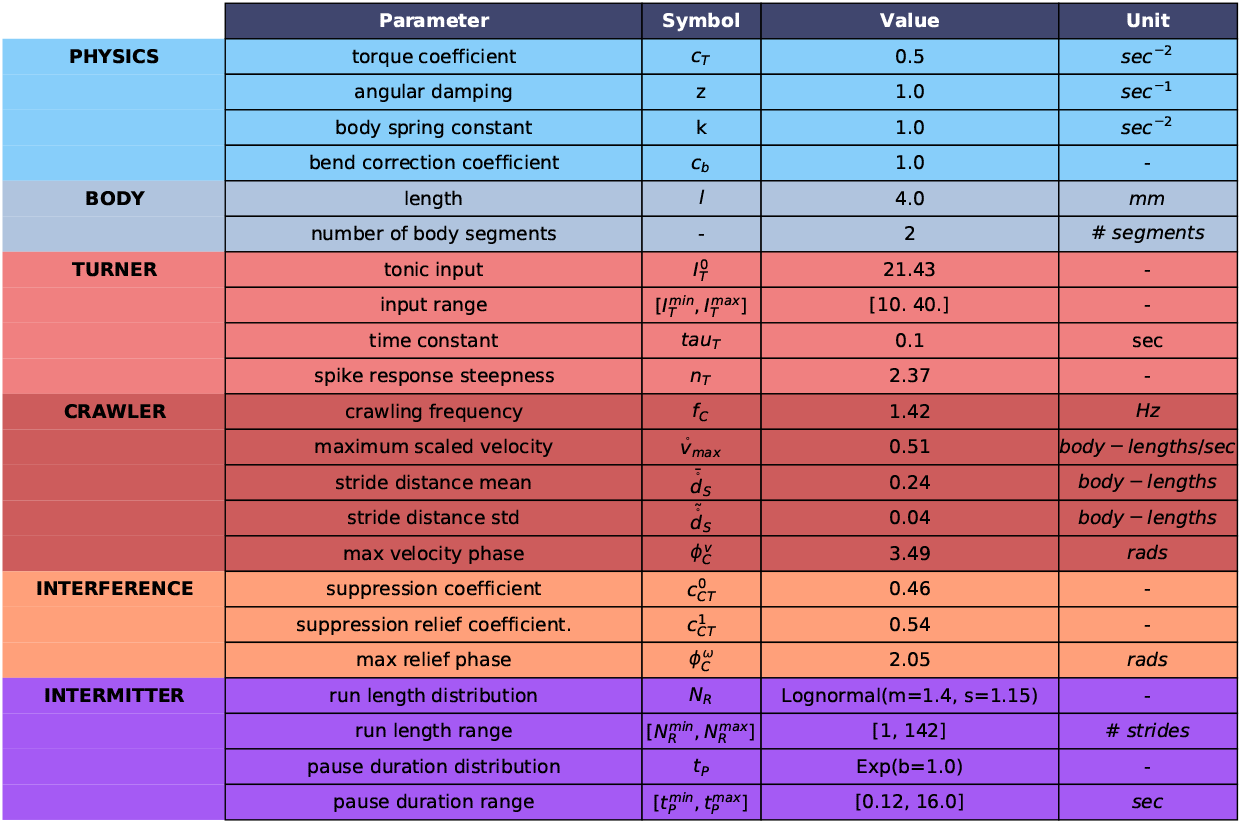
Locomotory model configuration. The parameters of the calibrated average locomotory model, organized per module.

**Video 3. Locomotory model for Drosophila larva**. The function of the intermittent coupled-oscillator locomotory model in ***Figure 1****B* is illustrated by adding its 4 modules stepwise: (i) crawler only, (ii) turner only, (iii) crawler + turner uncoupled, (iv) crawler + turner coupled (stride-phase attenuation of bending), (v) crawler + turner uncoupled + crawl intermittency (runs/pauses), (vi) crawler + turner coupled + crawl intermittency (complete intermittent coupled-oscillator model). The inset indicates active modules per clip.

http://computational-systems-neuroscience.de/wp-content/uploads/2024/10/2.mp4

#### Bisegmental body

The virtual body provides the interface between the behavioral architecture and the simulated environment. Its motion in the plane is defined by a forward velocity *v* and an angular velocity ω, which together determine the displacement and orientation of the larva. Any locomotory model that dynamically generates these two quantities is sufficient for a point-body or single-segment implementation.

For the coupled-oscillator model proposed here, we adopt a bisegmental body consisting of a front and a rear segment connected by a single bending degree of freedom (see ***Video 4***). The posture of the body is therefore described by the bending angle *θ*_*b*_ between the two segments. Following ***Wystrach et al. (2016)***, lateral bending dynamics are modeled using a torsional analogue of a mass–spring–damper system. In the standard formulation, translational and torsional systems are defined as

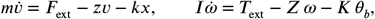

where *I* is the moment of inertia, *Z* the angular damping, and *K* the torsional stiffness resisting deformation.

For convenience, we express these dynamics in terms of angular acceleration per unit inertia by introducing the reduced coefficients

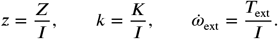

The external drive 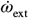 is generated by the turner module (described below). During crawling, angular motion is further modulated by a crawl-phase–dependent suppression coefficient *c*_*CT*_ (*t*) = *c*_*CT*_ (*ϕ*_*C*_ ) (see Crawler–turner coupling). Together, these contributions yield

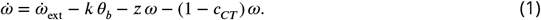

The original torsional model of ***Wystrach et al. (2016)*** omits two aspects of larval turning. First, it does not distinguish between bending velocity ω_*b*_ (change of *θ*_*b*_) and whole-body orientation velocity ω. Second, it does not correct the bending angle during forward motion. The bisegmental model resolves the first limitation by making ω and ω_*b*_ distinct variables. To address the second, we introduce a simple linear correction that gradually realigns the rear segment to the orientation of the front segment during forward displacement:

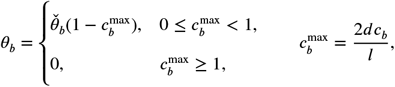

where *d* is the forward displacement during a timestep, *l* the body length, 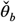 the bending angle before correction, and *c*_*b*_ = 1 the bend-correction coefficient.

Forward motion itself is generated independently by the crawler module, which directly supplies the instantaneous velocity *v* without additional biomechanical dynamics.

**Video 4. Bisegmental larva-body simplification**. The first panel shows the original tracked larval body with 12 midline points and a 22-point contour. Subsequent panels illustrate successive simplifications: removal of the contour, replacement with rectangular segment proxies, and finally a two-segment representation. The absolute head orientation angle *θ* is preserved, and the single bending angle *θ*_*b*_ is defined as 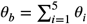 across the anterior body.

http://computational-systems-neuroscience.de/wp-content/uploads/2024/10/1.mp4

#### Crawler module

The crawler module generates forward velocity *v* under a tonic activation signal *I*_*C*_, which is supplied by the intermittency module. Specifically, during crawling epochs the intermittency module provides a constant tonic input *I*_*C*_ > 0, whereas during pauses it sets *I*_*C*_ = 0. The tonic input directly regulates crawling frequency, such that

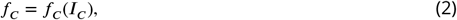

with *f*_*C*_ taking its empirically measured value when *I*_*C*_ is held constant during the short-duration simulations performed here.

Forward velocity generation depends on three parameters: the larval body-length *l*, the scaled displacement per stride 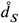, and the crawling frequency *f*_*C*_ . For each individual virtual larva, the parameters *l*, 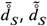, and the baseline *f*_C_ are assigned during initialization, where 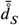 and 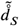 denote the mean and standard deviation of each larva’s stride displacement.

Crawling is modeled as an oscillatory process. Each oscillation cycle produces one forward stride of average length 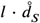, representing the net effect of a single peristaltic contraction cycle. After each stride a new 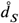 value is sampled from the empirically fitted distribution:

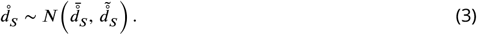

Across the oscillation, forward velocity follows a phase-dependent profile fitted to the average empirical stride waveform (see ***Appendix 2***–***Figure 1****B*). The resulting generative equation for in-stantaneous forward velocity is

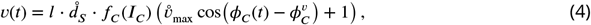

where *ϕ*_*C*_ (*t*) is the current phase of the crawler oscillator, 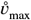 is the maximal scaled velocity, and 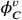 the phase at which this maximum occurs.

Thus, the intermittency module controls whether the crawler oscillates (run) or remains inactive (pause), while *f*_*C*_ (*I*_*C*_ ) and 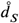 jointly determine the stride-by-stride forward progression of the virtual larva.

### Turner module

The turner module generates a torque-like lateral bending command *A*_*T*_ (*t*) under a continuous activation input *I*_*T*_ (*t*). This non-dimensional output is scaled to an external angular acceleration by the coefficient *c*_*T*_ (in s^−2^) and applied to the body as a bending drive (see Eq. 1):

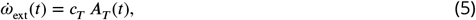

which produces the instantaneous angular velocity ω(*t*) and angular acceleration 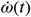 of the larva’s lateral body motion.

#### Oscillatory process

Previous work has proposed that larval turning is driven by an underlying oscillatory process generating alternating left–right bends (***Wystrach et al., 2016***). We confirmed the presence of such a slow rhythm empirically by detecting a dominant frequency of approximately *f*_*T*_ ≈ 0.4 Hz in the angular velocity time series of freely moving larvae. Motivated by this observation, we model the turner as a lateral oscillator extending the framework introduced in ***Wystrach et al. (2016)***.

The oscillator consists of two mutually inhibiting units, a left component (*E*_*L*_, *C*_*L*_) and a right component (*E*_*R*_, *C*_*R*_), where each pair includes an excitatory neuron *E*_*x*_ and an inhibitory neuron *C*_*x*_. The instantaneous firing response of each neuron to a cumulative input *x* is given by:

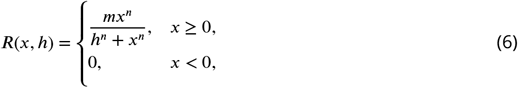

where *h* denotes the half-response threshold.

#### Neuromodulatory control

The tonic activation input *I*_*T*_ (*t*) excites both sides of the oscillator and modulates its gain and adaptation dynamics. Specifically, *I*_*T*_ (*t*) alters the gain *g*(*I*_*T*_ ) and the adaptation time constant τ_*H*_ (*I*_*T*_ ) of each neuron’s adaptation variable *H*_*x*_ according to:

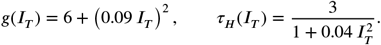

The resulting dynamics of the left-side excitatory and inhibitory neurons are:

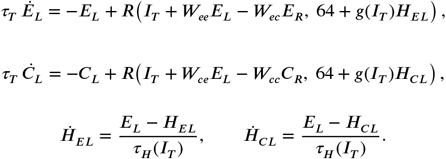

(The dynamics of the right-side neurons are defined analogously.)

Mutual inhibition between left and right components drives the system into stable antiphase oscillations, while the adaptation variables ensure periodic switching between sides. Under the baseline input 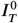 the oscillator generates alternating bending with dominant frequency *f*_*T*_ . Perturbations of *I*_*T*_ (*t*) modulate both the amplitude and frequency of the oscillations, which allows the turner to express context-dependent behaviors such as weathervaning and head casting.

The instantaneous lateral bending command is defined as the difference in excitatory activity:

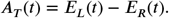

#### Parameter calibration

Achieving realistic angular motions requires joint calibration of the turner parameters and the biomechanical body parameters—namely, the angular damping *z*, the bend deformation resistance *k*, and the torque scaling coefficient *c*_*T*_ . The complete calibration procedure is provided in ***Appendix 2***.

#### Crawler–turner coupling

To capture the empirically observed interaction between forward crawling and lateral bending, we couple the crawler and turner oscillators by imposing a crawler-phase–dependent attenuation *c*_*CT*_ (*ϕ*_*C*_ ) on the angular velocity ω (see Eq. 1). During each stride cycle, the turning drive is suppressed by a baseline attenuation coefficient 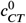, reflecting the mechanical constraint exerted by the ongoing peristaltic wave. Our kinematic analysis shows that this suppression is partially relieved during a specific phase interval of the stride, corresponding to moments when the anterior body is maximally free to bend laterally.

The degree of additional relief is governed by the relief coefficient 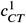, and the transition between full suppression and partial relief can be implemented in two alternative ways. In *square* mode an abrupt transition, where maximum relief is applied only within a fixed phase window 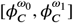 of the stride. In *phase* mode a smooth Gaussian kernel centered at phase 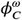, representing gradual lifting and reinstatement of attenuation around the phase of maximal bending freedom.

These two alternatives are implemented through the attenuation factor

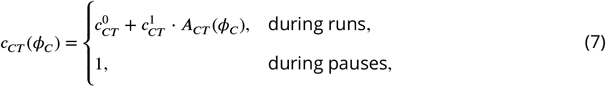

where the relief profile *A*_*CT*_ (*ϕ*_*C*_ ) is defined as

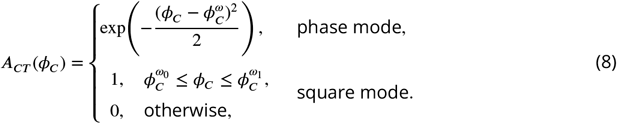

During pauses, the attenuation is removed (*c*_*CT*_ = 1), reflecting the empirical observation that large-amplitude head casts occur only when the larva is not executing a stride and thus the cranial body segments are free to bend without interference from peristalsis.

The coupling parameters 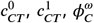 (phase mode) or 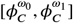 (square mode) are optimized to reproduce the experimentally measured average angular displacement as a function of stride phase. The full calibration procedure is described in ***Appendix 2***, and the resulting optimal Gaussian kernel is shown in ***Figure 3***.

#### Intermittency module

The intermittency module governs transitions between runs and pauses and serves as a place-holder for any mechanistic or statistical model capable of generating such switching dynamics (***Figure 1****B*). In the present implementation, the module provides the tonic input *I*_*C*_ to the crawler module: during runs, *I*_*C*_ is held at a positive constant value that maintains oscillatory crawling, whereas during pauses the module sets *I*_*C*_ = 0, immediately suppressing forward velocity (*v* = 0) and allowing large-amplitude head casts to occur.

For this study, run–pause switching is modeled statistically. At each transition, the duration of the upcoming state is sampled from empirical distributions fitted to experimental data. Pause durations are naturally measured in time units. For runs, we employ an equivalent metric by expressing run duration in terms of the number of consecutive crawling strides (“stridechains”), which reflects the stride-based structure of larval locomotion.

The fitted distributions for both pause durations and stridechain lengths are provided in ***Table 1***. These statistics fully determine the state durations used in the intermittency module for the simulations reported here.

#### Odorscape and olfactory sensor

Olfaction is introduced at the intermediate layer of the behavioral architecture to enable chemotactic behavior. The virtual larva carries a single olfactory sensor at its anterior tip; thus, both reorientation and forward displacement immediately affect the sensory input it receives. Odor sources are represented as Gaussian concentration fields *C*(*r*), where *r* denotes the distance from the odor center, the concentration peaks at *C*_0_ (in *μ*M), and spreads with standard deviation σ:

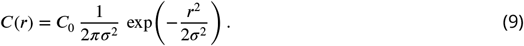

Following ***Gomez-Marin et al. (2011)***, we assume that olfactory perception responds to relative changes in odor concentration according to a Weber–Fechner–like relationship: increases or decreases in concentration modulate the perceptual variable *A*_O_ in proportion to Ċ/*C*. In addition, *A*_O_ relaxes back toward zero with a slow decay, yielding the dynamical equation

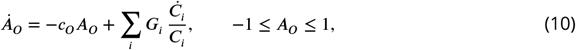

where *c*_O_ = 1 is the decay coefficient, *C*_*i*_ is the instantaneous concentration of odor *i* at the larval sensor, and *G*_*i*_ is its odor-specific gain.

To translate olfactory perception into motor modulation, we adopt the mechanism proposed by ***Wystrach et al. (2016)*** (***Figure 5***). Here, *A*_O_ perturbs the turner activation *I*_*T*_ away from its baseline value 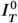 within the admissible range 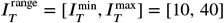:

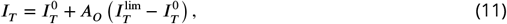

with

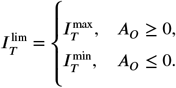

Positive *A*_O_ values thus enhance turning (approach), while negative values suppress it (avoidance), providing a unified modulation rule for attractive and aversive odors.

#### Mushroom body module

The mushroom body (MB) module implements the spiking network model of the larval olfactory pathway and MB published by ***Jürgensen et al. (2024)***. It consists of conductance-based leaky integrate-and-fire neurons and includes 21 olfactory receptor neurons (ORNs), 21 projection neurons (PNs), 21 local interneurons (LNs), 72 Kenyon cells (KCs), the anterior paired lateral neuron (APL), two mushroom body output neurons (MBONs), and two dopaminergic neurons (DANs) mediating reward and punishment. ORNs connect one-to-one to PNs and LNs; PNs provide sparse, random excitatory input to the KCs, while the APL provides global inhibitory feedback. All KCs excite both MBONs; each MBON in turn drives excitatory or inhibitory feedback onto the corresponding DAN.

Learning occurs at the KC::MBON synapses and is driven by a two-factor plasticity rule. A presynaptic KC spike sets an exponentially decaying eligibility trace *e*_*i*_(*t*) at synapse *i*, defining a window in which reinforcement can influence plasticity. When a DAN spike occurs, it provides a reinforcement signal *R*(*t*) that reduces synaptic strength proportionally to the current eligibility:

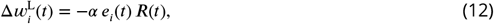

where α > 0 is the learning rate. The eligibility trace follows:

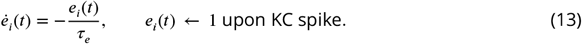

To balance learning-induced synaptic depression and to implement experimentally observed forgetting during odor-only trials, each KC::MBON synapse also undergoes homeostatic plasticity:

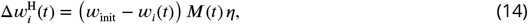

where *M*(*t*) is the postsynaptic MBON activity and η is the homeostatic rate. The total weight update at each integration step is:

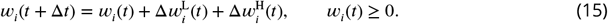

The MBON outputs encode the larva’s learned odor valence. We compute it from the spike counts of the approach-encoding MBON_+_ and avoidance-encoding MBON_−_ (***Owald et al., 2015; Owald and Waddell, 2015***):

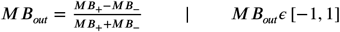

This output directly provides the modulatory gain used by the motor layer to bias orientation and chemotactic steering following learning.

## Acknowledgments

This project received funding from the Ministry of Culture and Science of the State of Northrhine Westphalia through the science network ‘iBehave’ (https://ibehave.nrw/) within the program “Netzwerke 2021”, and from the German Research Foundation through the Research Unit ‘Structure, Plasticity and Behavioral Function of the *Drosophila* mushroom body’ (DFG-FOR 2705, grant no. 365082554, https://www.uni-goettingen.de/en/601524.html). PS received a stipendship within the Research Training Group ‘Neural Circuit Analysis’ (DFG-RTG 1960, grant no. 233886668). We would like to thank Michael Schleyer and Juliane Thoener for providing the tracking dataset and for valuable discussions on the data analyses. We thank Bertram Gerber for valuable discussion and Anna Morozova for substantial support with the generation of the video and figure material.

## Appendix 1

### Metric definition

#### Body length

The instantaneous body-length of an individual larva fluctuates during crawling due to subsequent stretching and contraction. Its histogram is well fitted by a Gaussian distribution (data not shown). Therefore individual larva length *l* is defined as the median of the midline length across time (total length of the line connecting all 12 midline points). All spatial parameters including displacement and velocity can be scaled to this body-length, converting spatial units *m* or *mm* to dimensionless body-length units. Scaled spatial metrics are denoted by an additional° over the metric symbol.

#### Segmentation and angular metrics

To specify the body segmentation providing the most suitable contact/rotation point for the definition of the bending ω_*b*_ and orientation ω angular velocities we analyse their relationship in a subset of 40 larvae. Tracking of 12 midline points allows computation of the absolute orientation of 11 body-segments and the respective 10 angles *θ*_1−10_ between successive body segments (***Figure 2****F*). We define *θ* as the head-segment absolute orientation in reference to the x-axis because this defines the movement orientation of the animal. We ask how ω results from the bending of the body as this is captured by the 10 angular velocities ω_1−10_. The regression analysis depicted in ***Figure 1****B* shows as expected that ω depends primarily on the front angular velocities while this dependence decays as we move towards the rear segments, in line with previous studies (***Lahiri et al., 2011***). Timeshift analysis also shows that the front 3 angles change concurrently while angles further down the midline are increasingly lagging behind (data not shown). The correlation analysis depicted in ***Figure 1****C* shows that the sum of the front 5 angular velocities best correlates to ω. In other words the cumulative body bend of the front 5 segments best predicts head reorientation. Therefore we define the reorientation-relevant bending angle as 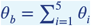 (***Figure 2****F*). The remaining 5 angles between the rear body-segments can safely be neglected as they do not contribute to reorientation. This analysis results in a segmentation of the body in a front and a rear segments of length ratio 5 : 6. The segmentation process is demonstrated in ***Video 4***.

#### Forward velocity

To define forward velocity we need to choose which midline-point is most suitable to track and which velocity metric to use for defining the start and end of a stride. To this end we perform stride annotation of 3-minute tracks of a population of 20 larvae using each of 24 candidate instantaneous velocity metrics, namely the velocities of the 12 points, the component velocities of the rearest 11 points parallel to their front segment’s absolute orientation and finally the centroid velocity. To compare the candidate metrics we compute the spatial *cv*_*s*_ and temporal *cv*_*t*_ coefficient of variation of the annotated strides for each larva to assess how variant their time duration and displacement is. We finally compute the mean 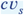 and 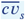 across individuals. In ***Figure 1****A* the spatiotemporal stride variance is shown for each candidate metric. We choose the metric that provides the minimal spatial and temporal stride variance, assuming that strides of an individual larva are more or less stereotypical in both duration and displacement (***Heckscher et al., 2012***). Our study reveals that the centroid velocity is the most suitable metric for stride annotation. All spatial metrics are therefore computed via this point’s displacement.

**Appendix 1—figure 1.**
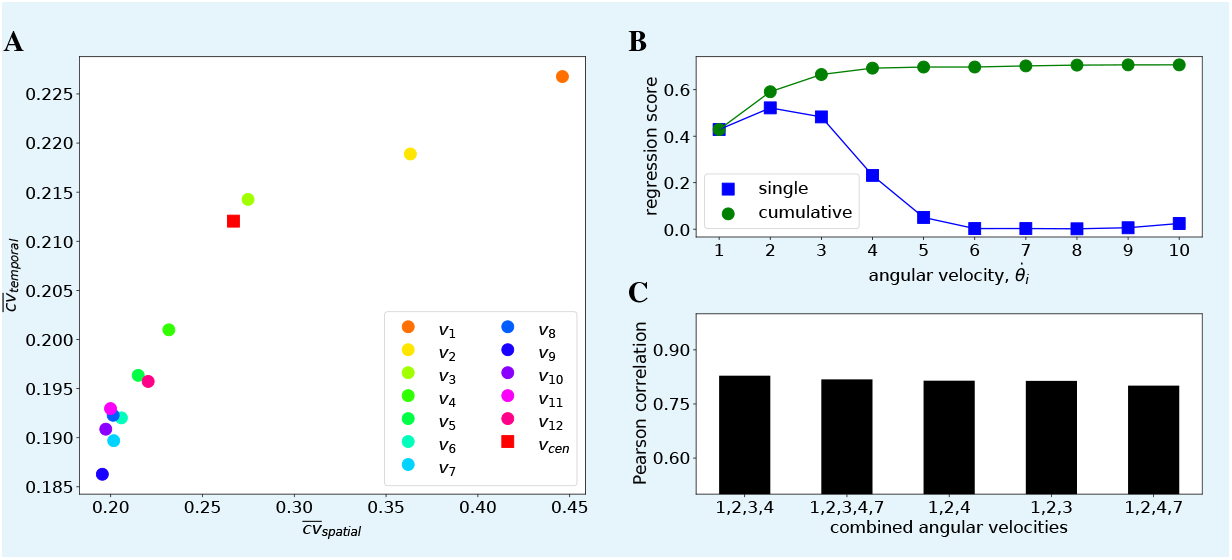
Segmentation and velocity definition. **A:** Forward velocity definition. 13 candidate velocity metrics are compared for use in stride annotation of 3-minute tracks of a population of 30 larvae. For each candidate the mean coefficient of variation of temporal duration 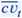 and spatial displacement 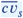 of the annotated strides is shown. Midline point 9 velocity provides the most temporally and spatially stereotypical strides, therefore it is selected as the reference forward velocity for stride annotation and model fitting. *v*_*cen*_ : centroid velocity, *v*_1_ − *v*_12_ : 1^*st*^-12^*th*^ point’s velocity. **B:** Regression analysis of individual and cumulative angular velocities ω_*i*=1−10_ to orientation angular velocity ω. When considered individually, ω_2_ best predicts reorientation with the ω_3_ and ω_1_ following. When considered cumulatively the anterior 4 ω_*i*_ allow optimal prediction of reorientation velocity. **C:** Correlation analysis of the sum of all possible ω_*i*_ combinations to ω. The sum 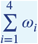 shows the highest correlation therefore we define 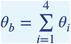 as shown in A. For illustration purposes only the 5 highest correlations are shown.

#### Track tortuosity

Track tortuosity is quantified by the straightness index (S.I), a metric previously used in larva-track analysis (***Sims et al., 2019***), computed by advancing a fixed time window along the track and calculating at each point the ratio of the straight line distance to the actual distance travelled. This index, which varies from 0 (no movement) to 1 (straight line movement), can capture very well the complexity of the movement at various scales (set by the window time frame) throughout the track. As the time window decreases the smoothing effect is also reduced revealing increasing track details. Different window sizes have been used ranging from 2 to 20 seconds to capture both large-scale spatial trajectories and small-scale local movements. Tortuosity was computed for each larva through time and revealed changes in movement as larva alternated between straight line relocation, changes of direction and different degrees of tortuosity. Hence, a change in the S.I. along a larva’s track captures the magnitude of the change in movement pattern from intensive, area-restricted searching movements (higher tortuosity) to extensive, straighter line movements (lower tortuosity), and vice versa, across a wide range of spatial scales.

#### Epoch annotation

Strides (*S*) are annotated using the scaled forward velocity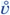. First we apply fourier analysis to detect the dominant crawling frequency within a suitable range 1 ⩽ *f*_*C*_ ⩽ 2.5 (***Figure 3****A*). From this we derive the reference stride duration 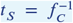. Then epochs are annotated under a number of constraints (***Figure 2****C*):

- Each stride is contained between two 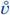 local minima.
- The 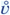 local maxima contained in the epoch needs to exceed a threshold : 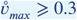.
- The duration of the epoch *t* needs to range within 0.75 ⩽ *tS*/⩽2. This allows individual strides to temporally vary without overlapping so that adjacent strides can be concatenated in stridechains.

After stride annotation the displacement due to each individual stride is computed for each larva and divided by the larva’s body-length (***Figure 2****A-C*). The individual distributions are well fitted by Gaussians (data not shown). Therefore 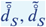 are defined as the average and standard deviation of the scaled displacement per stride for each larva.

Crawling runs (*R*) are defined as uninterrupted sequences of successive strides, also termed stridechains (***Figure 2****C*). Stridechain length is the number of concatenated strides in a run and is a discrete metric equivalent to the continuous crawling run duration. If the locomotory model under evaluation does not generate *v* oscillations and therefore strides can not be detected, runs are annotated using the plain forward velocity *v* timeseries as in a previous study (***Sakagiannis et al., 2020***). We first define a suitable threshold *v*_*thr*_ by detecting the minima of the pooled *v* bimodal distribution. Then we define runs as epochs where constantly *v* ⩾ *v*_*thr*_. Pauses are then defined as epochs containing no strides (or equivalently not overlapping with runs) and during which 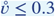 (or *v* ≤ *v*_*thr*_).

The turn epochs (*T* ) are contained between pairs of successive sign changes of the orientation angular velocity ω. Annotated epochs of the left (*T*_*L*_) and right (*T*_*R*_) turns yield the respective amplitudes of the turn-angle 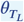 and 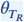 as the absolute total change in the orientation angle *θ* (***Figure 2****G*), which can then be pooled into the overall absolute amplitudes of the turn-angle *θ*_*T*_ . Notably by this definition turns include both headcasts (*H*) occurring during crawl-pauses and weathervaning (*W* ) occurring during runs. Turns occurring exclusively during runs and pauses yield the turn-angle amplitudes of weathervaning *θ*_*W*_ and headcasts *θ*_*H*_, respectively.

## Appendix 2

### Model calibration

Here we describe in detail the calibration procedure for a locomotory model of an average idealized individual based on a reference dataset of 150 larvae tracked while freely exploring a Petri-dish over 3 minutes. The pipeline consists of 3 initial steps, which can be performed independently of each other and set the configuration parameters of the crawler, intermittency and turner modules respectively. In a final step that builds upon the results of the initial three, the parameters of the crawl-bend phase-locked interference are set. The first two steps are directly defined by the kinematic analysis results. The third and the final steps involve an optimization process to reach the parameter set best fitting the empirical data. Notably, along with the intrinsic parameters of a module, a number of additional module-related parameters might need to be defined, such as the body-length for the crawler module and the angular-motion relevant physical parameters for the turner module.

**Appendix 2—figure 1.**
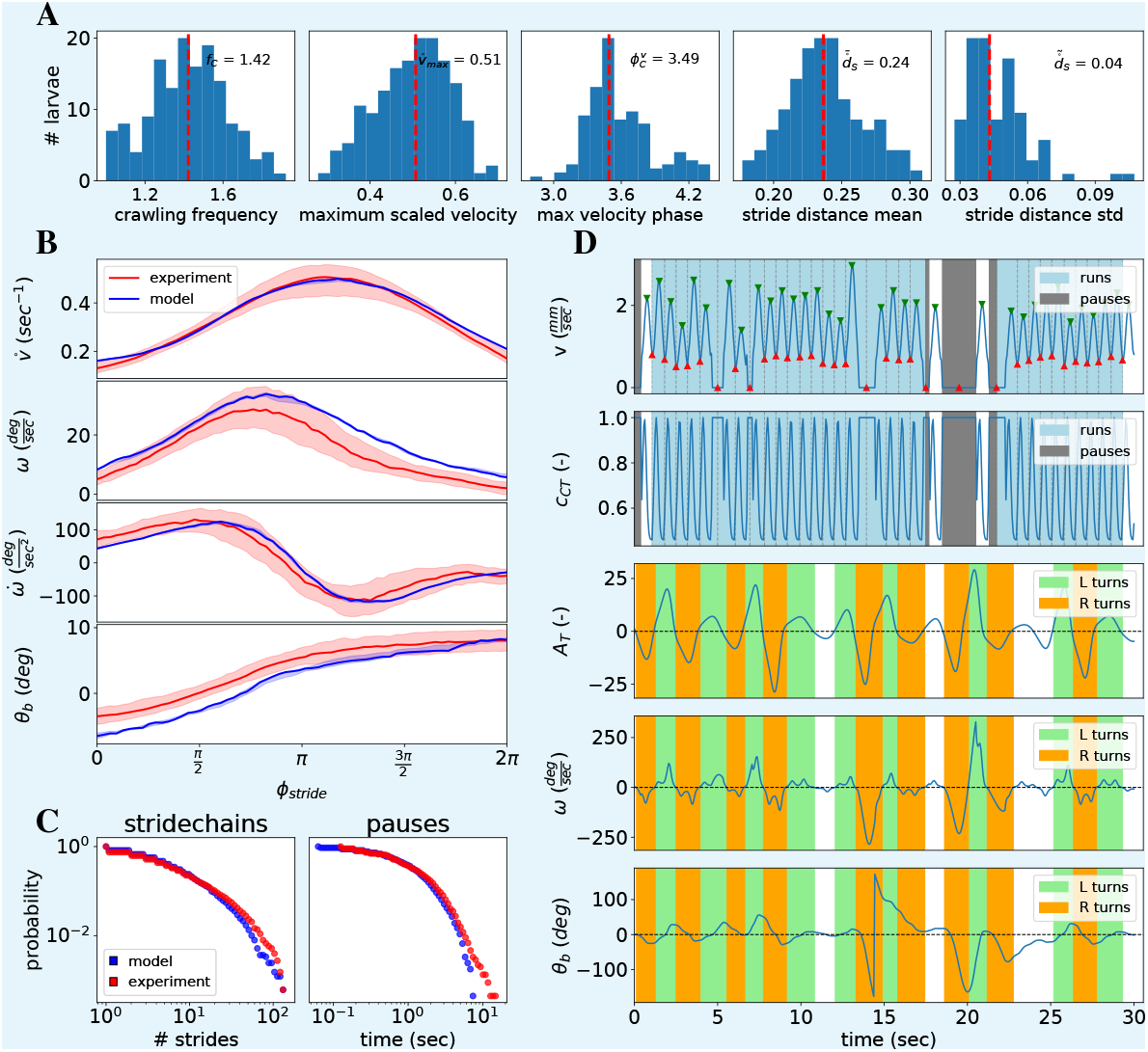
Average locomotory model summary. **A:** Distribution of stride-cycle related parameters over the empirical dataset. Red dotted line denotes the median value used in the Crawler module of the average locomotory model. **B:** Normalized average curves of angular metrics during the stride cycle for each individual larva. Red line denotes the group median. **C:** Pooled distribution of runs and pauses over the entire dataset (blue). Runs are detected as chains of concatenated strides (stridechains). Stridechains and pauses generated by the fitted distributions are shown in red. These distributions are used in the Intermitter module of the average locomotory model. **D:** Sample simulation of the model with all modules active. The model features additionally bend correction due to forward motion and crawling phase-coupled suppression of angular motion.

For the hereby described average-larva model no noise is introduced to any module input or output and no parameter variability is allowed across the population of the generated virtual larvae. The sole source of stochasticity then is due to the random distribution sampling processes operated during the behavioral simulations. For the analysis of the generated datasets the exact same pipeline is used as for the empirical dataset. The configuration of the final locomotory model is illustrated in ***Table 1***.

#### Forward motion

For the Crawler module a set of 5 parameters need to be defined, along with a default body-length. This parameter-set is adequate to dynamically generate realistic forward velocity *v* oscillations as defined in Eq. 4. All parameters are set to the median value across the empirical larva group. In ***Figure 1****A* the empirical distributions of the 5 crawler-module parameters are shown along with the median value selected for the average module.

#### Crawl intermittency

For the Intermittency module the sampling distributions for the run and pause epochs need to be defined. For both a continuous time-duration distribution is define while for the former an additional discrete distribution of number of concatenated strides per stridechain (run) is computed. In all cases, the pooled distribution of the given epoch across the entire dataset is approximated by power-law, exponential, Levy, log-normal and combined log-normal/power-law distributions. All distributions are truncated within an empirically observed range and are fully defined by their usual arguments. The best-fitting candidates from each class are sorted according to their Kolmogorov-Smirnov distance *D*_*KS*_ from the empirical distribution and the one exhibiting the minimum *D*_*KS*_ is selected for use in the model’s intermittency module (***Table 1***). A validation is shown in ***Figure 1****D* where the distributions of the generated epochs (blue) match the empirical ones (red).

#### Angular motion

Realistically calibrating the locomotory model in the angular domain is more demanding. The Turner module only generates a torque-equivalent output while the torsional-spring body model applies both angular damping and restorative force due to the body-bend angle (Eq. 1). We need to calibrate the 3 involved physics parameters, namely the angular damping *z* in *sec*^−1^, the bend deformation resistance *k* in *sec*^−2^ and the torque coefficient *c*_*T*_ in *sec*^−2^. Additionally the parameters of the turner module itself need to be specified. Here we study two implementations. The first is the above described neural oscillator, for which we need to define the baseline tonic input 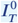, the spike-response steepness coefficient *n*_*T*_ and the time constant τ_*T*_ . The second is a simple sinusoidal oscillator, for which we need to define the baseline amplitude 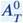 and frequency *f*_*T*_ . In order to have an empirical reference for isolated angular-only motion while avoiding the effect of crawling on the angular motion and we select the detected pause epochs exceeding 3 seconds and derive the pooled distributions for 3 angular metrics, namely the bending angle *θ*_*b*_, the angular velocity ω and the angular acceleration 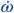 . We run an optimization algorithm aiming to minimize the Kolmogorov-Smirnov distance between the simulated and empirical distributions for these 3 angular metrics. In each iteration a Turner module of a given configuration is simulated for 3 minutes while Eq. 1 is applied to derive *θ*_*b*_ and ω. The obtained optimal turner and physics parameters are shown in ***Table 1***.

#### Crawl-bend interference

For the crawl-phase dependent attenuation of angular motion we study two implementations, each having three parameters that define the attenuation coefficient *c*_*CT*_ (*ϕ*_*C*_ ) during the stride-cycle (Eq. 7). To achieve this we use a genetic algorithm (GA) optimization process to reach a best-fitting parameter-set for this module. Although optimization is only allowed to search the parameter space for these three parameters, each configuration is evaluated using a complete virtual larva model, combined with the optimal configurations of all other modules defined in the previous three calibration steps. During each iteration of the GA process a group of 30 model configurations is simulated for 3 minutes in free exploration conditions. The best 6 configurations are then mutated and/or combined in a novel generation. The evaluation function minimized is two-fold.Firstly the distributions of the three angular metrics derived from the simulated track, namely the bending angle *θ*_*b*_, the angular velocity ω and the angular acceleration 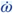 are compared to the empirically measured target distributions via the Kolmogorov-Smirnov distance as described above for the angular motion. Secondly, stride-cycle analysis is carried out for each simulated larva. Detected strides are interpolated in over *n* = 64 bins in a 2π cycle and direction-normalized by inverting the right-turning strides. The average stride-cycle curve 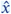 is then computed for each of the three metrics and compared to the average empirically measured curve *x* via the evaluation metric defined below. The metric has been selected so that the different scales of angles, angular velocities and accelerations are normalized before being summed. The final evaluation metric to be minimized is defined as the cumulative error across the three angular metrics for both the distributions and the average stride-cycle curves.

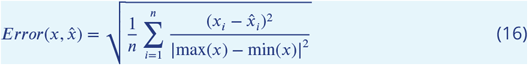

Here we study two configurations for the crawl phase-dependent suppression of the angular motion, namely a smooth gaussian relief curve and an acute square transition as described in Eq. 7.The target empirical and optimal model’s average curve for the angular velocity ω are shown in ***Figure 1****B*.

A sample simulation of the complete model is shown in ***Figure 1****D*. The Crawler-generated forward velocity oscillations (1st row) attenuate the angular motion in a phase-locked manner (2nd row) resulting in low angular velocity during runs and acute headcasts during pauses (3rd row). The bending angle is additionally restored to 0 during forward motion (4th row).

## Notes

http://computational-systems-neuroscience.de/wp-content/uploads/2024/10/1.mp4

http://computational-systems-neuroscience.de/wp-content/uploads/2024/10/2.mp4

http://computational-systems-neuroscience.de/wp-content/uploads/2024/10/3.mp4

http://computational-systems-neuroscience.de/wp-content/uploads/2024/10/4.mp4

http://computational-systems-neuroscience.de/wp-content/uploads/2024/10/5.mp4

http://computational-systems-neuroscience.de/wp-content/uploads/2024/10/6.mp4

http://computational-systems-neuroscience.de/wp-content/uploads/2024/10/7.mp4

http://computational-systems-neuroscience.de/wp-content/uploads/2024/10/8.mp4

http://computational-systems-neuroscience.de/wp-content/uploads/2024/10/9.mp4

### Competing Interest Statement

The authors have declared no competing interest.

### Summary of Updates

This version of the manuscript has been revised to update the following: In response to the elife public reviews, we substantially revised the manuscript to improve clarity, methodological transparency, and alignment between empirical data and model implementation. Major conceptual concerns were addressed by expanding the description of the behavioral architecture and explicitly motivating our coarse-grained, modular approach. We clarified that the goal is not to provide a fully neuromechanical account, but a flexible framework that integrates models at different abstraction levels and can incorporate more detailed mechanistic modules in future work. The Model section was significantly restructured. We moved essential explanations from the Appendix into the main text, added explicit formulations of the locomotory and intermittency modules, and clarified the rationale behind selecting a simplified bisegmental body representation. We expanded the description of the crawl-bend interference, the empirical basis for its phase-dependent attenuation, and its role in unifying head-casting and weathervaning. A new subsection, Limitations of the study, was added and rewritten for clarity, consolidating limitations raised by the reviewers and explicitly acknowledging the absence of segmental neuromechanics, explicit neural switching, handedness, long-horizon exploration, and social interactions. The Exploration results now clearly define dispersal and pathlength metrics, explain why only short-horizon dynamics are assessed, and discuss how these relate to MSD and to long-term data such as Sims et al. (2020). We additionally reference studies on handedness and clarify inter-individual variability. The Chemotaxis section was rewritten to provide a transparent, step-by-step description of how odor concentration modulates turning amplitude and frequency, how weathervaning and head-casts emerge from the same oscillator mechanism, and why the implementation should be viewed as a qualitative proof-of-concept rather than a quantitatively calibrated model. For Associative learning, we clarified the experimental paradigm (CS+ vs. neutral odor), the role of the mushroom body (MB) model, and the sequential, open-loop coupling between learning and behavior. We explain how MB output is transformed into odor-gain parameters and how population-level preference indices correspond to empirical group assays. Figures were reordered and corrected throughout, captions expanded, and the text revised for consistency, readability, and alignment with reviewer feedback. Finally, the underlying larvaworld software package was updated following reviewer-reported issues, and dependency pinning was relaxed to improve robustness.

